# Proteasome complexes experience profound structural and functional rearrangements throughout mammalian spermatogenesis

**DOI:** 10.1101/2021.06.18.447862

**Authors:** Dušan Živković, Angelique Sanchez Dafun, Thomas Menneteau, Adrien Schahl, Sandrine Lise, Christine Kervarrec, Ana Toste Rêgo, Paula C. A. da Fonseca, Matthieu Chavent, Charles Pineau, Odile Burlet-Schiltz, Julien Marcoux, Marie-Pierre Bousquet

## Abstract

During spermatogenesis, spermatogonia undergo a series of mitotic and meiotic divisions on their path to spermatozoa. To achieve this, a succession of processes requiring high proteolytic activity are in part orchestrated by the proteasome. The spermatoproteasome (s20S) is specific to the developing gametes, in which the gamete-specific α4s subunit replaces the α4 isoform found in the constitutive proteasome (c20S). Although the s20S is conserved across species and was shown to be crucial for germ cell development, its mechanism, function and structure remain incompletely characterized. Here, we used advanced mass spectrometry (MS) methods to map the composition of proteasome complexes and their interactomes throughout spermatogenesis. We observed that the s20S becomes highly activated as germ cells enter meiosis, mainly through a particularly extensive 19S activation and, to a lesser extent, PA200 binding. Additionally, the proteasome population shifts from c20S (98%) to s20S (>82-92%) during differentiation, presumably due to the shift from α4 to α4s expression. We demonstrated that s20S, but not c20S, interacts with components of the meiotic synaptonemal complex, where it may localize via association with the PI31 adaptor protein. In vitro, s20S preferentially binds to 19S, and displays higher trypsin- and chymotrypsin-like activities, both with and without PA200 activation. Moreover, using MS methods to monitor protein dynamics, we identified significant differences in domain flexibility between α4 and α4s. We propose that these differences induced by α4s incorporation result in significant changes in the way the s20S interacts with its partners, and dictate its role in germ cell differentiation.

## Introduction

Spermatogenesis is a process of cell differentiation, whereby a part of the population of stem cells called spermatogonia (SPG) enter the differentiation pathway and develop into spermatids, which then enter the spermiation process to become fully developed spermatozoa. Spermiogenesis is the differentiation of haploid round spermatids into elongated spermatids. During this maturation, a number of cell remodeling events are completed, including several critical cell-specific remodeling processes such as DNA condensation, mitochondrial reorganization, production of the flagellum and cytoplasm removal (1). SPG first divide mitotically and develop into spermatocytes (SPC), which enter meiosis. Meiosis requires duplication of the genetic material, its condensation, recombination between the maternal and paternal homologue chromosomes and re-distribution of the recombined chromosomes into separate cells. This is immediately followed by another division resulting in daughter cells called spermatids (SPT) with only one recombined copy of each chromosome. This entire process is facilitated by the Sertoli cells (SER) that serve to support developing gametes (2). Recombination is a particularly lengthy process requiring the formation of special bridges across paired chromosomes, called synaptonemal complexes (SCs), which enable the exchange of parts of chromosomes (3). Overall, spermatogenesis and spermiogenesis are both proteolysis-intensive processes with high protein turnover that require intense engagement of the primary proteolytic machinery of the cell, the proteasome (4).

The proteasome is a macromolecular proteolytic machinery in charge of controlled degradation of proteins (5). The proteasome core, also known as the 20S proteasome, is a symmetric, barrel-like structure that consists of a catalytic chamber formed by β subunits, which is sealed on both ends by a ring of α subunits (5–7). The 20S associates with various proteins and protein complexes, the most notable being the proteasome activator complexes 19S regulatory particle (to form the 26S proteasome), PA28αβ, PA28γ and PA200. These regulators modify the 20S activity and substrate specificity in order to regulate various processes such as cell division, differentiation, heat shock response, DNA repair, immune response, apoptosis, and many others (8–15). Moreover, the proteasome complexity is further increased by the presence of several proteasome subtypes where one or several subunits of the constitutive complex are replaced by alternative isoforms. Thus far, in addition to the constitutive 20S proteasome (c20S), immuno-(i20S), thymo-(t20S), and spermato-proteasomes (s20S) have been identified and described in the literature (16).

The s20S is a proteasome subtype in which the standard α4 subunit (PSMA7 gene) is replaced by α4s (PSMA8 gene) that is expressed exclusively in gamete cells (17). Previous studies in PSMA8^-/-^ KO mouse models have established that the α4s subunit is essential for spermatogenesis (14, 18). Functionally, mammalian s20S has been shown to degrade acetylated histones via association with PA200 and play a role in DNA damage repair in spermatocytes and the maturation of spermatids (11, 14, 19). Double strand break (DSB) repair was shown to be dependent on s20S-mediated degradation of non-histone substrates (19, 20). Cell cycle-mediating proteins were also reported as substrates of s20S (21, 22). However, despite these reports, s20S remains understudied and there is currently a lack on basic information on the exact nature and relative stoichiometry of the regulators binding to the s20S during spermatogenesis. In this context, revealing the dynamics of the 20S proteasome composition and partners throughout spermatogenesis could help explain how the s20S functions are conveyed, and further elucidate the underlying molecular mechanisms.

Here we use advanced mass spectrometry (MS)-based approaches to map interactomes of mammalian s20S during male germ differentiation and compare them with those of c20S. Our analyses in whole testes and in isolated male germ cells revealed specific s20S partners and profound changes in the dominant 20S proteasome subtypes present throughout differentiation. Incorporation of α4s into s20S was strikingly correlated with an increased association with 19S, PA200 and PI31 regulators, and with an overall proteasome activation, especially regarding its trypsin- and chymotrypsin-like activities. Moreover, using hydrogen-deuterium exchange (HDX)-MS and molecular dynamics simulations we identified conformational differences between α4 and α4s, providing a molecular rationale for the observed differences between c20S and s20S.

## Results

### Spermatoproteasome represents a major fraction of total 20S proteasome in mammalian testes

To examine differential expression patterns of s20S and c20S, as well as other proteasome-associated factors, we analyzed tissue-specific human proteome maps (17) and observed that the expression pattern of the PA200 and 19S regulators closely follows that of α4s (**Fig. S1A**). We noted that both male and female gonads express s20S (**Fig. S1A**), but given that the only reported role for s20S is in male germ cells we decided to focus our analysis on whole testes and purified male germ cells. This does not exclude that the s20S may play a still unknown role in females.

To map the relative abundance of s20S, we used freshly frozen bovine testis lysate where the s20S is highly expressed. We immunopurified (IP-ed) 20S proteasome using the MCP21 antibody that recognizes the α2 proteasome subunit (present in all 20S proteasome subtypes), as described previously (23). The 20S proteasome was then further separated from its regulators by an additional Size Exclusion Chromatography (SEC) step (24). After IP and SEC, the purified 20S proteasomes were analyzed on a LC-MS system, using a previously optimized method (25). We were able to identify not only the proteasome subunits that are conserved among the different 20S subtypes, but also the specific catalytic subunits of the c20S (β1, β2, β5) and the i20S (β1i, β2i, β5i), as well as the testes-specific α4s subunit of the s20S (**Fig. 1A**; **Table S1**). Taking the β5 subunit abundance as reference, the results show that most of the 20S proteasomes in testis contain exclusively the three constitutive catalytic subunits (64% ± 1 %), while some complexes with different combinations of constitutive and immune subunits are also present (26, 27) (**Table S1)**. Additionally, we could quantify the proportion of s20S using the MS signals of α4 *vs*. α4s subunits, by assuming that they have similar ionization yields based on their high sequence identities (**Fig. S1B**). In this way we established that the s20S constitutes 48 ± 1 % of the total testis immunopurified proteasomes, which will thereafter be called “testes 20S pool”.

**Figure 1.**
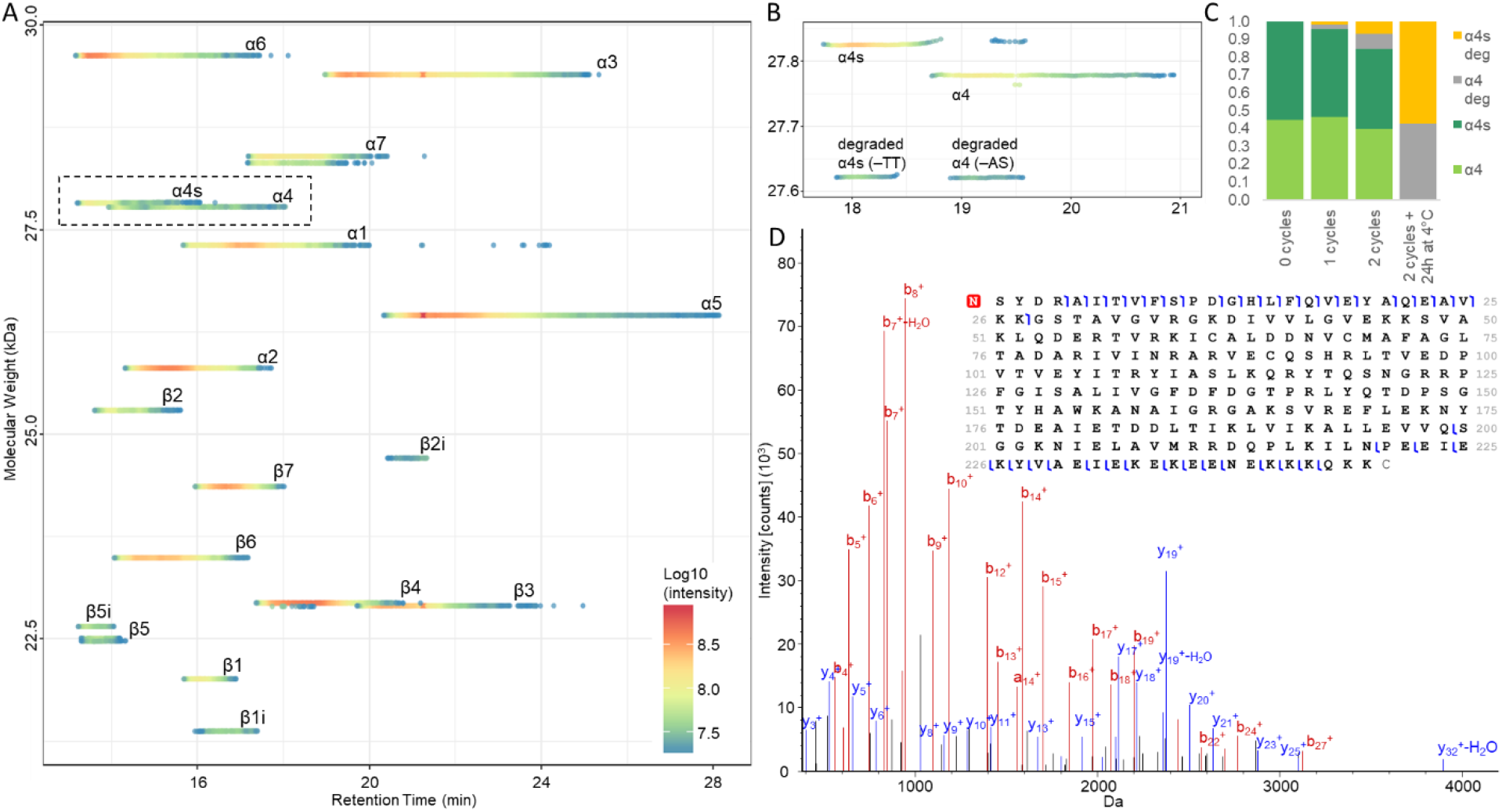
**(A)** 3D proteoform footprint of the LC-MS trace of 20S subunits immunopurified from bovine testis, generated with VisioProt-MS (28). The names of the c20S subunits are indicated next to the corresponding signal. **(B)** Close-up view of the 3D proteoform footprint of 20S subunits immunopurified from bovine testes after repeated freeze-thaw cycles, resulting in truncated proteoforms of α4 and α4s. **(C)** The truncated α4 and α4s to the full-length proteins ratio increase with repeated freeze-thaw cycles. **(D)** Top-down LC-MS/MS of α4 identifies the degradation as the loss of the last two amino-acids.

In addition to quantifying different proteasome species, we were able to observe various PTMs (**Table S2**), that were in accordance with published data (25, 29, 30). Interestingly, we noticed that α4s harbors the same PTMs as the α4 proteoform, *i.e*. loss of the initial methionine and N-terminal acetylation. Additionally, we observed that repeated freeze-thaw cycles led to formation of truncated versions of α4 and α4s subunits that lack the last two amino acid residues (**Fig. 1B–C**), which were later confirmed by LC-MS/MS Top-Down (TD) sequencing (**Fig. 1D**). A similar loss in mass, corresponding to a truncated version of α4, was recently described in rat and rabbit (30). Given that our results indicate that these lighter α4/α4s isoforms are produced upon storage, most likely via proteolytic cleavage of their solvent accessible C-termini, we consider them unlikely to be biologically relevant. We also noted +73 Da and −13 Da mass differences observed for β6 and β2, respectively (**Fig. S2**). TD sequencing confirmed that in our sample these subunits differ from the official (curated) Swiss-Prot entries by 1 amino-acid (G233E for β6 and N252T for β2) but are in agreement with predicted TrEMBL entries. These mutations are barely visible using bottom-up proteomics where proteins are commonly identified with only a few peptides, illustrating the benefits of TD-MS for proper proteoform characterization and database curation. Taken together, our TD-MS analysis of the immunopurified proteasome pool from bovine testes revealed that s20S represents a major fraction of the total proteasome, suggesting a key functional role.

### The s20S has its specific set of interactors in bovine testes

To further characterize the s20S in gamete cells, we mapped its interactome using an IP strategy. We performed two IPs from bovine testes lysates using either an anti-α4s-specific antibody generated according to (31), and validated as shown in **Fig. S3**), which targets only the s20S, or an anti-α2 antibody, which recognizes all proteasome types, and compared the relative abundances of co-immunopurified proteins. In a first set of experiments, lists of proteasome interacting proteins (PIPs) specific to α2 and α4s were obtained by comparing the anti-α2 or anti-α4s IPs against a control IP (antibody directed against rat CD8-OX8, as used previously (32)). 3867 proteins were validated and quantified in the anti-α2 and anti-α4s IPs, with a total of 1177 proteins exhibiting over a 2-fold change (FC) enrichment (significance threshold at *p<0.05*) compared to the control (**Fig. S4, S5**). To highlight putative specific or enriched components in c20S and s20S interactomes, we compared the relative abundances of each of these proteins in the anti-α2 IP (total 20S) *vs*. anti-α4s IP (s20S). To do so, protein abundances were normalized based on the 20S content in each IP (estimated as the average of the abundances of the non-catalytic 20S subunits), then the fold changes (anti-α4s IP / anti-α2 IP) and their significance (*p*-value) were calculated and represented as a Volcano plot (**Fig. 2A**). Here an unequal distribution of enriched proteins is observed, skewed towards the anti-α4s IP. One possible explanation for this observation is that the anti-α4s antibody might fetch 20S interactors that appear below the detection threshold when analyzing the whole 20S interactome (*i.e*., with the anti-α2 antibody only). However, we also acknowledge that the anti-α4s polyclonal antibodies used may display more non-specific bindings, compared to monoclonal antibodies used in the anti-α2 IP. We therefore looked specifically at proteins that might be of interest, namely proteasome- and ubiquitin-related proteins.

**Figure 2.**
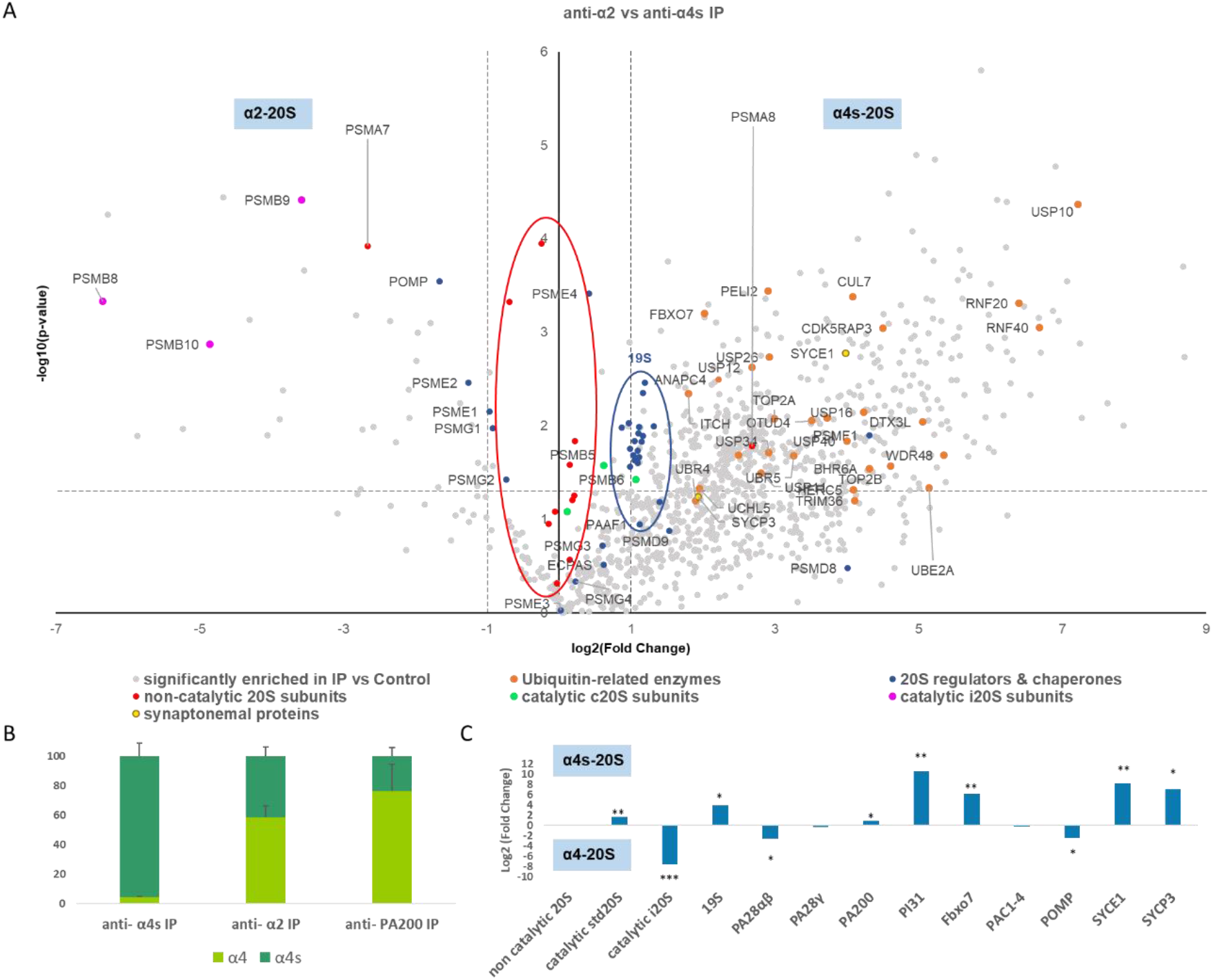
**(A)** Analysis of proteasome composition and interacting proteins in bovine testes. Distribution of enriched proteins between two IPs against anti-α2 (left hand side) and anti-α4s (right hand side). Only proteins that were enriched (FC>2, *p-value* <0.05) in the anti-α2 or anti-α4s IPs compared to the control IPs are displayed here. For clarity, gene names are used as protein identifiers. 20S non-catalytic and 19S subunits are circled in red and blue, respectively. **(B)** Estimated percentages of α4 and α4s in the different IPs. The protein intensities were normalized based on the 20S content and the relative abundances of α4 and α4s estimated for the different IPs. **(C)** FCs of 20S-associated regulators in the s20S (α4s-containing proteasome) relative to α4-containing proteasomes. The relative quantities of α4 and α4s in the anti-α4s and anti-α2 IPs enables to estimate the fraction of 20S-associated regulator in the α4s- and α4-containing proteasomes. Finally, FCs in α4s-*vs*. α4-containing proteasomes were calculated, as detailed in the experimental section. Graphics represent mean and standard deviations. All values are the means of three independent experiments. Stars indicate significance. *: p<0.05; **: p<0.01; ***: p<0.001.

The distribution displayed on **Fig. 2A** highlights the specific composition features of c20S and s20S complexes. Only proteins that were enriched (FC>2, *p-value* <0.05) in anti-α2 or in anti-α4s IPs compared to the control IPs are displayed in this comparison. We observed that the non-catalytic 20S subunits (except α4 and α4s - PSMA7 and PSMA8 genes, respectively) are found tightly centered around the y-axis (mean FC = 1.0 +/− 0.2), showing that the normalization of the dataset is well done and that there are no major variations in their relative abundance. As expected, the bait of the anti-α4s IP is found on the right (α4s protein - PSMA8, FC = 6.4, *p-value* = 0.016) whereas the subunit replacing it in the c20S is found on the left (α4 protein - PSMA7 gene, FC = 0.16, *p-value* = 12.E-4). We then used the relative abundances of the α4 (PSMA7) and α4s proteins (PSMA8) in the anti-α2 IP to estimate the part of s20S in the total pool of 20S proteasome in bovine testis (42 ± 7%; **Fig. 2B**), which is in good agreement with the results obtained by the TD approach. On the other hand, the anti-α4s IP contains 95 ± 9% of s20S (**Fig. 2B**), confirming the exclusive nature of the α4s subunit incorporation into the s20S (14, 18). Taken together, our data provide evidence that the α4s and α4 subunits do not co-exist within a hybrid 20S proteasome. This subunit exclusivity is in agreement with what has been previously observed for some i20S catalytic subunits (33, 34), and could result from exclusive expression of either the α4 or the α4s or from chaperone-mediated preferential incorporation of α4s in 20S proteasome.

We observed that the three i20S-specific catalytic subunits (PSMB8-10 proteins, **Fig. 2A**) were more abundant in the anti-α2 IP and were practically absent from the anti-α4s IP, implying that little or no immuno-subunits constitute the s20S (FC of immuno subunits in anti-α4s IP = 0.006, *p-value* = 7.E-5, **Fig. 2C**), as previously observed using an orthogonal immunodetection approach (18). This interesting result is also in agreement with the 6.1 fold decrease, in the anti-α4s IP, of PA28αβ (*p-value* = 0.01), known to interact preferentially with the i20S (23). Moreover, while the four 20S Proteasome Assembly Chaperones (PAC) are equally distributed in the α4- and α4s-containing 20S (**Fig.2A, Table S3**), the POMP maturation protein is 5.3 fold increased in the α4-containing 20S (*p-value* = 0.02, **Fig.2C, Table S3**). POMP is known to preferentially promote the assembly of i20S catalytic subunits over c20S ones (33, 35, 36), which agrees with our observation that immuno-catalytic subunits are low abundant in the anti-α4s IP.

Our analysis of main 20S interactors revealed that all subunits of the 19S particle could be quantified and their very tight distribution in the Volcano plot (*i.e*. very close FCs and *p-values*, **Fig. 2A**) emphasizes the quality of this dataset. Interestingly, we could measure an average increase of 2.13 ± 0.15 of 19S subunits in the anti-α4s IP compared to the anti-α2 IP. Since the anti-α4s antibody purifies almost exclusively α4s-containing 20S (95 ± 9%) and the anti-α2 IP contains a mix of proteasomes (42% of α4s-containing 20S), we can estimate that in whole testes, the abundance of the 19S bound to the s20S is approximately 15 times higher than the one associated with the c20S (*p-value* = 0.02) (**Fig. 2C** and **Table S3**). Interestingly, other known 19S interactors were also found to be increased in the anti-α4s IP compared to the anti-α2 IP. This is the case for the two main 19S-associated deubiquitinases USP14 and UCHL5 (FC = 3.2 and 3.9, *p*-values = 0.02 and 0.05, respectively), ADRM1 protein, a known 19S receptor of polyubiquitin chains (FC = 1.8, *p*-value = 0.01), as well as RPS27, the main cellular precursor of ubiquitin (FC = 1.9, *p*-value = 0.03). The K48 polyubiquitin chains identified by the LIFAGK(GG)QLEDGR peptide (Døskeland, 2006) were increased by a factor of 3.0 (*p*-value = 0.001) in the anti-α4s IP compared to the anti-α2 IP. Altogether this dataset suggests a higher loading of polyubiquitinated substrates onto s20S compared to total proteasome complexes. Accordingly, among the 40 ubiquitin-related enzymes (conjugating enzymes, ligases and deubiquitinylases) that are found significantly regulated between the two IPs, all were increased in the s20S interactome except TRIM21, an E3 ligase that is mainly involved in immune response (**Table S4**).

Another PIP found to be highly regulated in this dataset is PI31, a 20S interactor of controversial function (37). PI31 displayed a 20-fold enrichment in the anti-α4s IP compared to the anti-α2 IP (**Fig. 2A**, *p-value* = 0.01) and seems to be a preferred partner of the s20S (**Fig. 2C** and **Table S3**). Strikingly, a known heterodimerization partner of PI31, Fbxo7 (38), is also significantly more abundant in the s20S IP (FC 72; *p-value* = 0.007). Recent KO studies of Fbxo7 and Nutcracker, its heterolog in drosophila, indicate important roles of both proteins during spermatogenesis (39, 40). Due to the established role of PA200 in spermatogenesis (11), a complementary IP directed against PA200 was conducted and also showed that PI31 and Fbxo7 proteins were highly enriched in the anti-α4s IP compared to the PA200 IP (FCs = 12.2 and 4.1, *p-values* = 1E-04 and 0.05, respectively, **Fig. S6, S7**), suggesting that these proteins are not bound to the s20S through an interaction with PA200.

We found that the synaptonemal proteins SYCP3 and SYCE1 are specifically enriched in the anti-α2 and anti-α4s IPs, compared to their respective control IP (**Fig. S4, S5**), and significantly more abundant in the s20S interactome compared to the other α4-containing 20S isoforms (FCs = 294 and 3.8, respectively; *p-values* = 8E-03 and 0.05, respectively (**Fig. 2A, 2C** and **Table S3**). Previous imaging studies have suggested that these proteins interact directly with the s20S (18), and our dataset further demonstrates such specific interactions of the s20S with the SC.

Despite their proposed role in spermatogenesis (41), the nuclear PA28γ and PA200 regulators of the 20S core were either not (FC = 0.8; *p-value* = 0.4) or only slightly, but still significantly (FC = 2.0; *p-value* = 0.03), enriched in the s20S compared to the α4-containing 20S subtype (**Fig. 2C** and **Table S3**). Moreover, when comparing the relative abundances of α4 and α4s co-purified in the anti-α2 and in the anti-PA200 IPs, we observed similar values (**Fig. 2B**). Altogether, these results suggest that PA200 may not interact preferentially with the s20S, contrary to what was previously speculated (16). However, if the α4 and α4s isoforms are expressed in different cell types, as previously reported (14), they would not have to compete for PA200 association. Thus, higher association observed between PA200 and the s20S could be due to a higher PSMA8 gene expression in these cells.

Taken together, our IP strategy revealed that the s20S has an interactome distinct from that of the c20S. The s20S interactome we mapped here is enriched for the 19S particle, as well as different components of the UPS and synaptonemal proteins. The differences between the c20S and s20S interactomes suggest distinct functions of these proteasome variants.

### Proteasome dynamics analysis shows profound rearrangements throughout spermatogenesis

An important limitation of our whole testis analysis is the lack of information on the dynamic interactions that are frequently lost and/or rearranged upon tissue lysis (42), and information on proteasome dynamics throughout spermatogenesis. To provide a detailed map of the proteasome interactome during spermatogenesis, we initiated a study of individual germ cell populations. We harvested testes from young adult or pre-pubescent rats, and immediately dissected and digested them into individual cells. Rat germ cells were purified and separated either by sedimentation at unit gravity (SPG) or centrifugal elutriation (SPC and SPT), in order to obtain highly enriched cell populations. SER were also purified, since they are support cells of the germ cell lineage and might carry relevant information. Proteasome complexes from these purified fractions of cells were *in vivo* crosslinked to maintain protein-protein interactions (43) and then co-immunopurified using the antibody directed against the α2 proteasome subunit (23). Purified complexes were digested with trypsin and the resulting peptides analyzed on a nanoLC-MS/MS system, followed by protein identification and relative quantification. In parallel, lysates of individual groups of cells were directly analyzed without the IP step to obtain complementary information on protein expression and to validate the cell purification protocol. Among the 5750 proteins validated and quantified in the lysates using label-free MS, a pool of 12 proteins were unambiguously identified as specific markers of each cell type (**Fig. S8**). Proteasome complexes immunopurified from the different groups of cells were then analyzed using the same approach to observe changes in 20S proteasome composition throughout spermatogenesis (**Fig. 3**). None of the three i20S catalytic subunits could be detected in the four different cell types studied, suggesting that the i20S subunits previously detected in the whole bovine testis sample are likely derived from infiltrated immune cells. Another striking observation was that the α4s isoform almost completely replaced α4 at the SPC and remained the main isoform throughout spermatogenesis (**Fig. 3A)**. This trend was also observed in cell lysates (**Fig. 3B**), and may be driven by changes in the protein expression or stability of the two isoforms.

**Figure 3.**
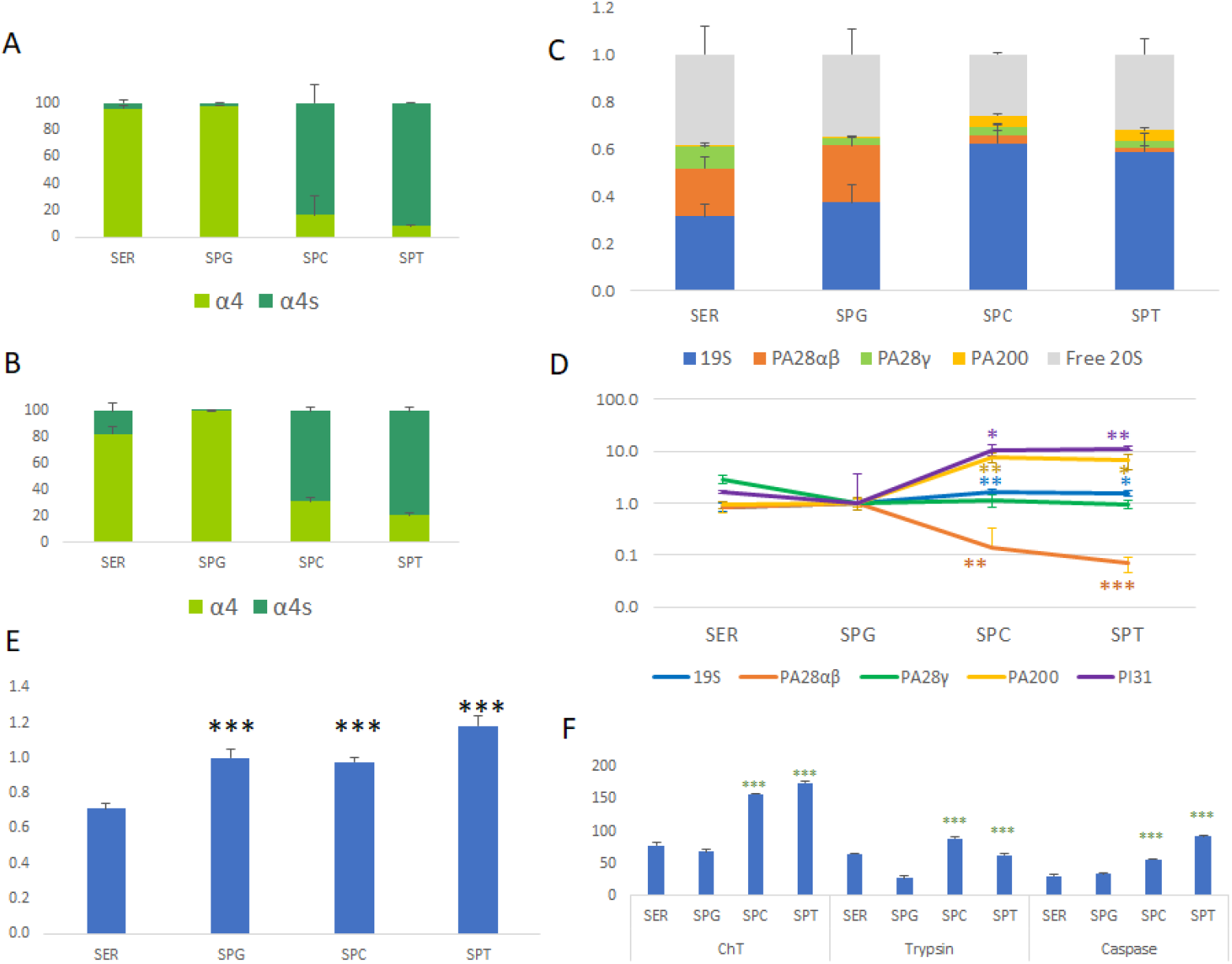
Composition and proteolytic activity of proteasome complexes purified from rat SER, SPG, SPC and SPT. Cells obtained from rat testes were separated and then analyzed using label free proteomics. **(A-B)** Proportions of α4 and α4s proteins in **(A)** immunopurified proteasome complexes and **(B)** cell lysates. **(C)** The distribution of the different proteasome complexes was estimated for each cell type by label-free MS (27). We considered the free 20S complexes and the ones associated with 19S, PA28αβ, PA28γ, and PA200 regulators. Error bars indicate standard deviation across three biological replicates. **(D)** Major changes in the composition of proteasome complexes throughout spermatogenesis. The variations of major 20S-associated regulators were measured throughout spermatogenesis using label-free quantitative proteomics. The SPG stage was used as a reference and stars indicate significance between SPC *vs*. SPG or SPT *vs*. SPG. **(E)** Changes in the amount of total 20S proteasomes throughout spermatogenesis. The variations of 20S proteasome subunits (all non-catalytic subunits except α4 and α4s) were measured throughout spermatogenesis using label-free quantitative proteomics and normalized with the MS signal of total protein. Stars indicate significance between SPG *vs*. SER or SPC *vs*. SER or SPT *vs*. SER. **(F)** Proteasome chymotrypsin-like (ChT), trypsin-like and caspase-like specific activities, expressed in pmol AMC released per minute and per mg of total protein, have been measured in the four cell types lysates throughout spermatogenesis. Stars indicate significance between SPC *vs*. SPG or SPT *vs*. SPG. Graphics represent mean and standard deviations. All values are the means of three independent experiments. *: p<0.05; **: p<0.01; ***: p<0.001.

Our analysis of proteasome activators (19S, PA28αβ, PA28γ, and PA200) indicated that the 19S is the predominant regulator associated with the 20S particles, whatever the cell type analyzed, and was bound from around 30% (in SER and SPG) up to 60% (in SPC and SPT) of the total 20S pool (**Fig. 3C**). The fraction of 20S-19S complexes thus significantly increases 1.7-fold (*p-value* = 0.01) when SPG proceeds into the SPC stage (**Fig. 3D**), and, as previously seen in bovine testes, the K48 polyubiquitin chains identified by the LIFAGK(GG)QLEDGR peptide were increased by a factor of 2.9 (*p*-value = 1E-03), highlighting a high demand for cellular ubiquitin-dependent proteolysis. While the nuclear PA28γ activator levels do not significantly change through spermatogenesis, its cytoplasmic counterpart PA28αβ significantly decreased (7.2-fold) from the SPG to SPC stages (*p-value* = 0.005) (**Fig. 3D**). The abundance of PA200 bound to the 20S core particle increases 7.5-fold in SPC (*p-value* = 0.002) and 6.6-fold in SPT (*p-value* = 0.01) compared to SPG, in agreement with the proposed role of PA200 in spermatogenesis (11, 44).

We also noted that PI31 has a dramatic increase (11-fold) in 20S core particle association in both SPC and SPT stages, compared to non-differentiated SPG cells (*p-values* of 2E-03 and 8E-04, respectively) (**Fig. 3D**). Accordingly, its known interactor Fbxo7 was only detected in proteasome complexes purified from SPC and SPT.

Given that 20S-associated regulators display different substrate specificities and subcellular localization (45), the observed variations in the panel of proteasome activators suggest profound changes in the biological function and/or subcellular localization of proteasomes throughout spermatogenesis. Overall, a large proportion of the 20S proteasome pool was bound to activators, reaching almost 75% in SPC cells, which represents a specific feature of testis cells as the proportion of activated 20S core is usually around 20 to 60% in other tissues (27). Moreover, all three germ cell types contain slightly but significantly higher amounts of total 20S compared to SER (**Fig. 3E**). Chymotrypsin, trypsin, and caspase-like proteolytic activities were significantly higher in meiotic and post-meiotic germ cells (SPC & SPT) compared to undifferentiated SPG (**Fig. 3F**), which is most probably the consequence of two major events, *i.e*. the complete replacement of α4 by α4s isoform and the increase in activator-associated 20S, in particular the higher loading of 19S and PA200 particles. No significant variation of 20S-associated regulators can be observed from SER to SPG cells which also display similar 20S proteasome composition, in particular their high content of α4. The observed variations in proteasome proteolytic activities and 20S-associated regulators in pre-meiotic (SPG) *vs*. meiotic and post-meiotic cell types (SPC & SPT) thus suggest either a possible increased ability of the s20S to interact with its activators, compared to the c20S, or a transcriptionally-driven increase in the quantities of these regulators in post-meiotic stages. To answer this crucial question, we analyzed variations of expression of these different proteins in the different cell lysates. Changes from SPG (containing exclusively c20S) to SPC and SPT (containing 85-90% s20S, **Fig. 3A**) were thus compared both in the immunopurified proteasome samples and in the lysates (**Fig. S9**).

Concerning α4s, its increase in lysates from the SPG to SPC stages is approx. two-fold higher than its increase in immunopurified proteasomes (*p-value* = 0.007, **Fig. S9**). This may be due to a chaperone-mediated resistance to α4s incorporation into functional 20S proteasomes, or to the lag time between synthesis of α4s and its incorporation in newly-assembled s20S. Increase in proteasome bound PA200 appears to be mainly the consequence of increased expression (**Fig. S9**), which is in accordance with the existing literature (19). However, given that we did not observe increased binding of PA200 to s20S in the whole testis interactome analysis, it remains difficult to draw conclusions about specific PA200 interactions at this stage. Finally, the increase in the association of 19S and PI31 with 20S proteasomes from SPG to SPC is, to a certain extent but not completely, explained by higher cellular abundances, since their increase in the immunopurified proteasomes are significantly higher than in full lysates (FC = 1.4 and 2.7; *p-value* = 0.03 and 0.006, respectively, **Fig. S9**).

Altogether this dataset indicates that changes in the composition and activation of 20S proteasomes from the SPC stage onwards may, at least partly, result from transcriptional regulation; however, at this stage we can’t exclude the possibility that other mechanisms and factors may be involved.

### Pull-Down assays show a preferential binding of the s20S to the 19S in vitro

In order to examine preferential interactions between α4s and PI31, 19S and PA200 (compared to α4), we took advantage of the testes 20S pool immunopurified in mg quantities from bovine testes. After incubation of these three partners of the 20S with either pure c20S (purified from bovine muscle) or the testes 20S pool (purified from bovine testes), we pulled-down complexes using anti-PI31, anti-19S or anti-PA200 antibodies-grafted beads, and estimated the relative amounts of α4 and α4s in the eluates using LC-MS/MS label free quantification. A control pull-down was run concurrently with the MCP21 antibody that equally recognizes the c20S and s20S. After normalization of the signal with the MCP21 IP, we found that PI31 was interacting more with proteasomes with α4 than with α4s **(Fig. 4A)** (*p-value* = 0.018). However, the 19S pull-down (anti-PSMC2) was much more efficient in co-purifying α4s than α4 (*p-value* = 0.0002) **(Fig. 4B)**. Finally, the anti-PA200 IP showed no significant difference between α4 and α4s pulldowns **(Fig. 4C)**. Altogether, these *in vitro* binary interaction experiments suggest that the c20S and s20S preferentially interact with PI31 and the 19S, respectively; while there is no clear evidence that PA200 interacts preferentially for any of these 20S subtypes.

**Figure 4.**
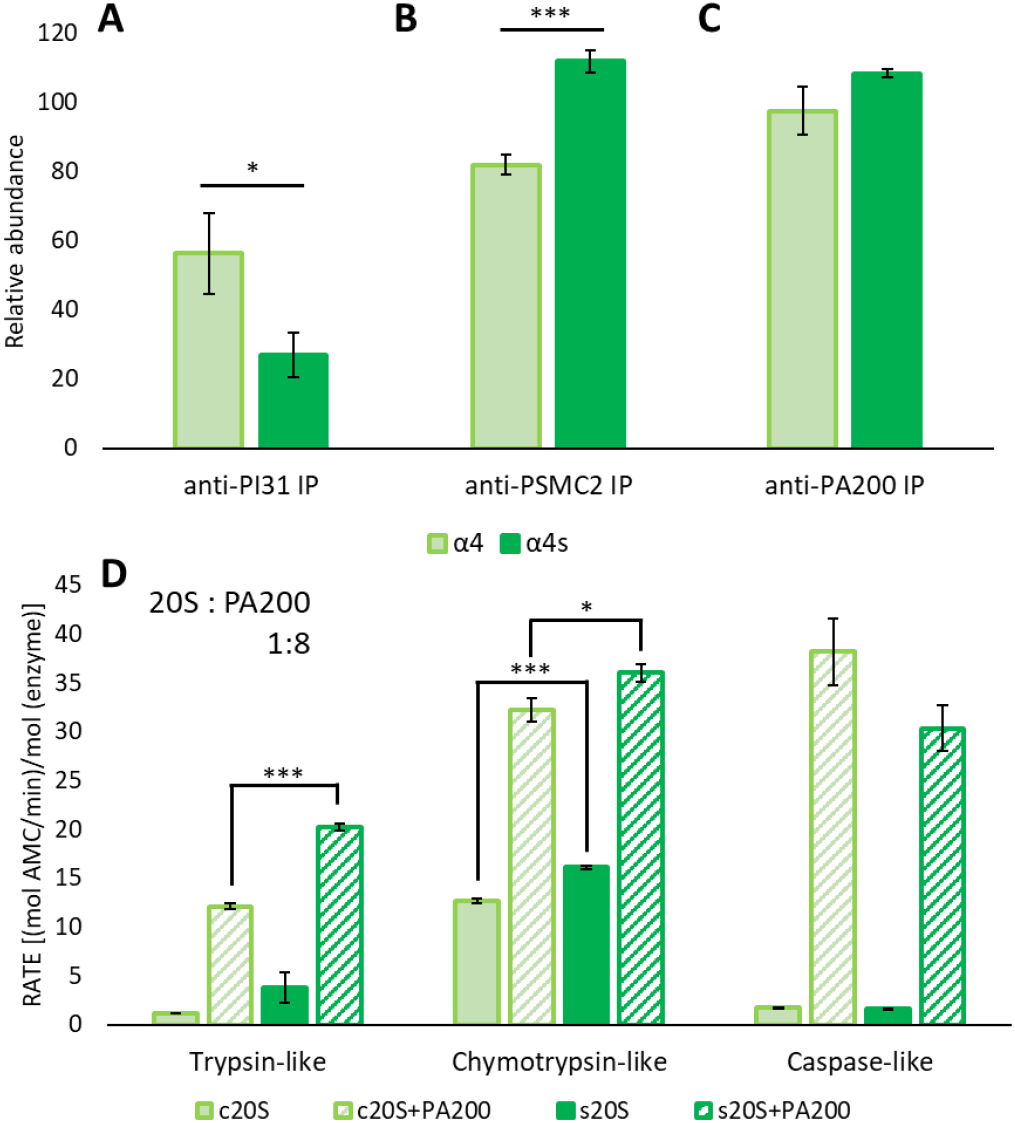
In vitro assays. Bovine c20S or testes 20S pool were incubated with **(A)** PI31, **(B)** the 19S or **(C)** PA200, and complexes were purified with sepharose beads grafted with their corresponding antibodies. After washes and elution, α4 and α4s subunits were identified and quantified by LC MS/MS label-free quantification and their abundances in anti-PI31 IP, anti-PSMC2 IP, and anti-PA200 IP were normalized with their abundances in the corresponding control IPs. Anti-α2 subunit control IP (for total 20S) was performed on the same reaction mix for each experiment. Graphics represent mean and standard deviations. All values are the means of three independent experiments. Stars indicate significance *: p<0.05; **: p<0.01; ***: p<0.001. The vertical axis represents the intensity of a protein in a pull-down experiment relative to its intensity in a control IP (in the case of 19S - average intensity of a set of 19S base proteins), when normalized for 20S abundance. **(D)** Comparison of three peptidase activities in proteasomes purified from bovine testis (s20S) or muscle (c20S). Activation of the proteasomes by PA200 was assayed at a PA200:20S molar ratio of 8:1. The bars are grouped by the peptide used to probe each activity: boc-LRR-AMC, suc-LLVY-AMC, and z-LLE-AMC, which probe the trypsin-, chymotrypsin- and caspase-like activities, respectively. Different complexes are represented by different colors. PA200 alone was tested as a control, and showed no intrinsic activity. 20S proteasome was used at a final concentration of 7.4 nM. All values are the means of three independent experiments. Stars indicate significance. *: p<0.05; **: p<0.01; ***: p<0.001.

### Testis and muscle proteasomes show distinct peptidase activities

We tested whether α4s incorporation into the 20S core induces any change in proteasome activity on its own or in complex with a regulator. We compared the testes 20S pool purified from bovine testes, which contains approximately 50% of s20S, and the c20S purified from bovine muscle in their ability to degrade fluorogenic substrates. The substrates assayed were boc-LRR-AMC, suc-LLVY-AMC and z-LLE-AMC, which probe the trypsin-like, chymotrypsin-like and caspase-like activity, respectively. The two proteasome samples were also activated with a PA200:20S molar ratio of 8:1. When comparing the basal activity of the proteasome, the s20S mix showed greater trypsin- and chymotrypsin-like activity, while the caspase-like activity was more or less the same (**Fig. 4D**). The addition of PA200 did activate the three protease activities of both 20S proteasomes. Although trypsin-like activity change upon PA200 binding is greater in c20S, the overall tryptic activation of the complex remains significantly greater in the s20S. The activated complex’s chymotrypsin-like activity is also greater in the s20S. The activation of the caspase-like activity is somewhat more pronounced in the c20S compared to the s20S. Overall, these results show that s20S seems to have higher trypsin and chymotrypsin-like activities than c20S, both at basal state and when activated with PA200.

### HDX-MS rationalizes the spermatoproteasome structural singularity

We recently implemented HDX-MS, a technique which reports on solvent accessibility and/or flexibility of the backbone amide protons, on the c20S and i20S proteasome complexes, in order to decipher their structural rearrangements upon replacement of their catalytic subunits and binding to PA28 regulators (46). Building on this expertise, we investigated the conformational differences due to the replacement of α4 by α4s. The relatively large amount (~180 μg per condition) of pure 20S required to perform such structural studies led us to use the testes 20S pool purified from bovine testes. This prevented us from comparing the deuteration of any peptides common to the c20S and s20S. However, 33 and 27 of the 53 and 47 peptides obtained upon pepsin digestion of α4 and α4s, respectively, were proteospecific (**Fig. S10**). We manually compared the deuteration heatmaps obtained for both α4 and α4s after alignment (**Fig. 5A**) and clearly identified two regions, encompassing residues 180-189 and 225-250, that were more readily deuterated in α4s compared to α4 (**Fig. 5A–C**). Interestingly, these two regions face each other on the outer surface of the α-ring, based on the c20S structure (**Fig. 5B**). To explore possible impacts of these differences on structure and dynamics of α4 and α4s, we employed 1 μs all-atom molecular dynamics simulations. We could detect ~64% more hydrogen bonds (H-bonds) between the NH groups of the backbone and water molecules for α4s than for α4 (**Fig. S11A**) in line with our HDX-MS results. Furthermore, the distribution of the crossing angle between the two last α-helices was broader in α4s compared to α4 (**Fig. S11B**, standard deviation = 8.3° and 6.2° for α4s and α4, respectively), indicating that the motion between these two helices is more important in α4s.

**Figure 5.**
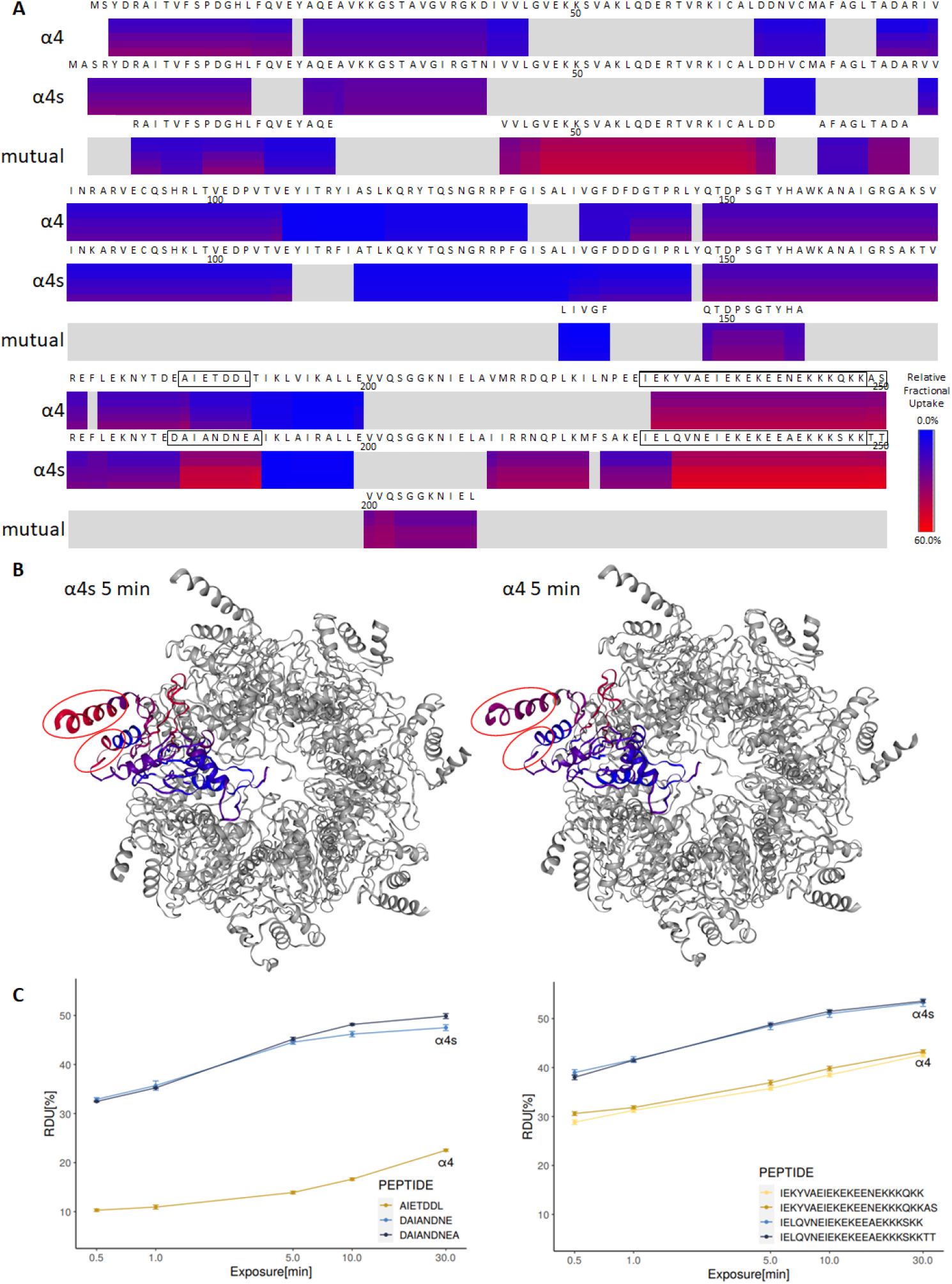
HDX-MS of the s20S *vs*. c20S. **(A)** Heatmap showing the relative deuterium uptake from 0% (blue) to 60% (red) of α4 and α4s at 0.5 min, 1 min, 5 min, 10 min and 30 min. **(B)** Cartoon representation of a homology model of α4s (left) and the structure of α4 (right) within the 20S bovine proteasome (PDB: 1IRU), viewed from the top of the α-ring. Residues of α4s and α4 are color coded with the relative deuterium uptake at 15 min. **(C)** Kinetics of deuteration obtained for 7 peptides of α4s or α4 encompassing residues 180-189 (left) and 225-250 (right), all showing a more limited deuteration of α4 compared to α4s.

Looking in more detail, we observed that these two helices were stabilized by H-bonds between three pairs of side chain residues (E235/K193, E231/K189 and Y228/D185 for α4 and E235/R193, E231/K189 and Q228/N185 for α4s) (**Fig. S11C**). Interestingly, the frequency of these H-bonds was found much higher in α4 than in α4s, especially between residues 235 and 189 (**Fig. S11, S12A**), explaining the higher motion of these helices, the larger number of H-bonds between its backbone and water molecules and the overall faster deuteration in α4s. In line with these results, the average Accessible Surface Area (ASA) of residues 184-198 and 225-241 was higher in α4s than in α4 (**Fig. S12B**). Altogether, our results indicate more dynamic movements of both 180-189 and 225-250 regions in the s20S compared to the c20S, potentially creating a different binding interface and resulting in the differences in the interactomes we mapped above.

## Discussion

Despite being recognized as important for mammalian male germ cell differentiation and spermatogenesis, the testis-specific proteasome isoform s20S remains incompletely characterized. Here, we combined state-of-the-art MS-based proteomic analyses with *in vitro* studies to conduct a comparison between s20S and c20S, using α4s and α4 as markers for each proteasome variant, respectively. We found that α4s represents ~42-45% of all α4 variants found in whole bovine testis, suggesting that the s20S is a major proteasome form in testicular tissue. We also showed that α4 and α4s do not co-exist within assembled proteasomes, in line with immunofluorescence imaging experiments (14). This exclusive incorporation of α4 or α4s subunits into proteasomes may be either due to a selective subunit incorporation or to a sudden change in protein expression levels. To address this question, as well as map dynamic proteasome changes that accompany spermatogenesis, we analyzed isolated populations of rat germ cells at different differentiation stages (SPG, SPC, SPT) and their supportive cells (SER). This demonstrated that SPG 20S proteasomes contained almost exclusively α4 (>98%), whereas pachytene SPC and SPT predominantly had α4s-containing proteasomes (>82%-92% α4s). This sudden change in the abundance of α4s vs. α4 may be driven by rapid changes in transcription or in protein turnover resulting in major differences in protein levels.

The shift of c20S to s20S observed from pre-meiotic SPG to meiotic SPC was accompanied by profound changes in the major 20S-associated regulators, in particular an increase in PA200 and 19S activators, concomitantly with a decrease in PA28αβ. Surprisingly, no change in PA28γ could be observed, although this 20S activator was previously shown to be relevant to male fertility (22, 41). It has been recently proposed that PA200 constitutes a major component of the s20S complex (11, 19) and that α4s may help the formation of PA200-capped complexes containing only standard catalytic subunits (19). Accordingly, our results indicate that PA200 is an important player in s20S function, as its level of incorporation into assembled proteasomes in SPC and SPT (around 8-10%) is at least 10-fold higher than in SPG and previously measured for several other cell types (27). However, our results also demonstrate that the 19S is the major activator of 20S core particles in germ cells, and in particular in meiotic SPC where α4s completely replaces α4. Indeed, the 19S is bound to around 30-40% of the total 20S pool in SER and undifferentiated germ cells and, strikingly, this proportion increases up to 60% in SPC and SPT cells, making meiotic and post-meiotic proteasomes the most activated proteasome complexes we have analyzed so far. This agrees with the increased proteolytic activities in SPC and SPT, compared to SPG and SER, in line with previous observations using whole testis lysate compared to muscle tissue (11). Another study also showed high levels of 19S-containing species in post-meiotic germ cells (18) and in whole testes (11, 19). Thus, contrary to what we expected from previous studies (11, 19), we clearly demonstrated that PA200 is far from being the sole activator of s20S during germ cell differentiation. The increased association between s20S and its 19S activator is probably aligned with high ubiquitin-dependent proteolytic requirements at the SPC stage, as suggested by increased polyubiquitin chains at the s20S complex in meiotic SPC compared to pre-meiotic SPG. In particular, meiosis I progression would require above all the ubiquitin-proteasome system (40, 47, 48), and, to a lesser extent, PA200 (14) which was shown to be rather involved in later acetylated histone turnover events (19).

We observed that PI31, previously described as an *in vitro* 20S proteasome inhibitor (37), but also a physiological 26S proteasome activator (40, 49) or assembly factor (50), together with its binding and stabilizing partner Fbxo7 (38), were both enriched in s20S interactomes compared to c20S but also PA200 IPs. Strikingly, both PI31 and Fbxo7 are essential for proper spermatogenesis in *Drosophila* (40) and mice (39), respectively. To our knowledge, our work is the first establishing a direct link between the α4s specific subunit of the s20S and the PI31/Fbxo7 axis. PI31 was shown to mediate proteasome transport in axons and dendrites in mice, by regulating the loading of proteasomes onto microtubule-dependent molecular motors (51). Accordingly, here we identified several microtubule-related proteins interacting with the s20S, which could facilitate such transport of s20S complexes. On the other hand, as a part of its E3 ligase activity, Fbxo7 targets proteins involved in cell cycle regulation (52). Thus, we speculate that PI31 and Fbxo7 might act as shuttle proteins for the s20S, targeting cyclins and other spermatogenesis-specific substrates; however, mechanistic details of s20S-PI31-Fbxo7 remain to be established.

Another class of s20S-specific interaction partners we identified are SYCE1 and SYCP3, components of synaptonemal complexes. These findings further support the proposed mechanism whereby s20S binds to the synapses along the axes of the meiotic chromosomes, to regulate this process through the degradation of specific proteins (18, 47). Although previous work suggested that SYCP3 could be a substrate of the s20S (18, 47), our data indicate high sequence coverages of SYCP3 and SYCE1 in our interactome study (25% and 34%, with 4 and 9 peptides, respectively), which is typical of true interactors. Moreover, recent work on s20S KO mouse models did not replicate the SYCP3 accumulation (14), further strengthening the argument for SYCP3 as a s20S interactor. Taken together, our quantitative interactome data indicate extensive differences between c20S and s20S, and show that spermatogenesis is accompanied by a major shift from c20S as the major proteolytic machinery towards s20S.

Given that the only difference between c20S and s20S is the presence of α4 or α4s isoforms, we employed HDX-MS to examine whether they display any differences in structure and dynamics. Our analysis revealed pronounced differences in flexibility of the C-terminal regions of the two isoforms, corresponding to the two last α-helices located on the outer, solvent-exposed side of both α4 and α4s structures. Our HDX-MS data showed that these two helices are more rigid (or stabilized) in α4 than in α4s. Molecular dynamics simulations provide a rationale for these observations, by predicting more stable hydrogen bonds between the two C-terminal helices in α4. In order to understand functional consequences of these differences, we performed pull-down assays with α4- and α4s-containing proteasome and immobilized proteasome regulators, PI31, PA200 and 19S. We observed differences in pull-down efficiency for PI31, which showed significantly greater affinity towards the α4-containing proteasome (c20S), and 19S, which had a higher affinity towards the s20S. However, we did not observe any differences in affinity of PA200. Finally, we also tested proteolytic activity of proteasomes purified from bovine testes (~50% s20S) and from the muscle (100% c20S) in two different conditions: alone and when activated by PA200. We measured that s20S has higher basal trypsin- and chymotrypsin-like activity, and this trend held upon PA200 activation. On the other hand, caspase-like activity for 20S alone was either the same or very similar, while the c20S-PA200 complex was more active than the s20S-PA200. Interestingly, upon PA200 binding, the fold-change in caspase-like activity is the most pronounced, confirming previous observations (53). Increased basal level of tryptic activity can lead to improved degradation of substrates that are highly positively charged, such as histones, in agreement with the report that showed histones to be the targets of the PA200-20S complex in the context of spermatogenesis (11). Overall, these activity measurements indicate significant difference in behavior of the α4s-containing proteasome, compared to the α4-containing proteasome.

Taken together, the large amount of data analyzed and presented in the context of this study highlight some key differences between c20S and s20S. Our results imply a more complex process of s20S regulation than previously suggested. Based on these, we can speculate that the structural differences between the s20S and c20S proteasome variants trigger the recruitment of specific partners and ubiquitin-related enzymatic modulators that are key for proteasome cellular relocalization to the SC and for the degradation of important meiotic players, respectively.

### Experimental Procedures

#### Proteome Repository Search

For data on relative expression at the protein level of the proteasome and proteasome-related genes we searched The Human Proteome Map portal (17) by querying the list of relevant genes.

#### Reagents

Unless stated otherwise, all reagents were purchased from Euromedex. The c20S, i20S, PA28γ and PA28αβ were purchased from Enzo Life Science. The mouse IgG1 anti-α2 antibody was produced from the hybridoma cell line MCP21 (European Collection of Cell Cultures).

#### Antibody development

Anti-α4s antibodies were produced by Biotem (Apprieu, France) using procedures described in SI Experimental Procedures.

#### Preparation of separated germ cells

Cells were obtained from rats using procedures described in SI Experimental Procedures.

#### LC-MSMS analysis

Bottom-up and top-down proteomics experiments were performed on an Orbitrap Fusion instrument coupled to an Ultimate 3000 chromatography system. Acquisition parameters and data analysis are detailed in SI Experimental Procedures.

#### HDX-MS and MD simulations

HDX-MS experiments were performed on a Synapt-G2Si (Waters Scientific, Manchester, UK) coupled to a Twin HTS PAL dispensing and labelling robot (Trajan Scientific, Milton Keynes, UK) via a NanoAcquity system with HDX technology (Waters, Manchester, UK), as described previously (46). More details, as well as MD simulation workflow can be found in SI Experimental Procedures.

#### In vitro assays

The activity assay, based on fluorogenic peptide degradation, and the pull-down assays are further detailed in SI Experimental Procedures.

#### Data availability

The mass spectrometry proteomics data have been deposited to the ProteomeXchange Consortium via the PRIDE (54) partner repository with the dataset identifier PXD027436.

## Acknowledgements

This work was supported by the French Ministry of Research (ANR-ProteasoRegMS to JM, Investissements d’Avenir Program, Proteomics French Infrastructure, ANR-10-INBS-08 to OB-S), University of Toulouse (grant to DZ), the Fonds Européens de Développement Régional Toulouse Métropole and the Région Midi-Pyrénées (OB-S). PCAdF and ATR were funded by the Medical Research Council (MC_UP_1201/5 grant to PCAdF).

## References

1. P. Calvel, A. D. Rolland, B. Jegou, C. Pineau, Testicular postgenomics: targeting the regulation of spermatogenesis. Philosophical Transactions of the Royal Society B: Biological Sciences 365, 1481–1500 (2010).

2. A.-F. Holstein, W. Schulze, M. Davidoff, Understanding spermatogenesis is a prerequisite for treatment. Reprod Biol Endocrinol 1, 107 (2003).

3. N. Hunter, Meiotic Recombination: The Essence of Heredity. Cold Spring Harb Perspect Biol 7, a016618 (2015).

4. C.-C. Hou, W.-X. Yang, New insights to the ubiquitin–proteasome pathway (UPP) mechanism during spermatogenesis. Mol Biol Rep 40, 3213–3230 (2013).

5. G. A. Collins, A. L. Goldberg, The Logic of the 26S Proteasome. Cell 169, 792–806 (2017).

6. M. P. Bousquet-Dubouch, B. Fabre, B. Monsarrat, O. Burlet-Schiltz, Proteomics to study the diversity and dynamics of proteasome complexes: From fundamentals to the clinic. Expert Review of Proteomics 8, 459–481 (2011).

7. I. Sahu, M. H. Glickman, Structural Insights into Substrate Recognition and Processing by the 20S Proteasome. Biomolecules 11, 148 (2021).

8. D. Aristizábal, V. Rivas, G. I. Cassab, F. Lledías, Heat stress reveals high molecular mass proteasomes in Arabidopsis thaliana suspension cells cultures. Plant Physiology and Biochemistry 140, 78–87 (2019).

9. P.-M. Kloetzel, The proteasome and MHC class I antigen processing. Biochimica et Biophysica Acta (BBA) - Molecular Cell Research 1695, 225–233 (2004).

10. M. Min, C. Lindon, Substrate targeting by the ubiquitin–proteasome system in mitosis. Seminars in Cell & Developmental Biology 23, 482–491 (2012).

11. M.-X. Qian, et al., Acetylation-Mediated Proteasomal Degradation of Core Histones during DNA Repair and Spermatogenesis. Cell 153, 1012–1024 (2013).

12. D. Sohn, et al., The Proteasome Is Required for Rapid Initiation of Death Receptor-Induced Apoptosis. Mol Cell Biol 26, 1967–1978 (2006).

13. C. E. Widjaja, et al., Proteasome activity regulates CD8+ T lymphocyte metabolism and fate specification (2017) https:/doi.org/10.1172/JCI90895 (September 14, 2021).

14. Q. Zhang, S.-Y. Ji, K. Busayavalasa, J. Shao, C. Yu, Meiosis I progression in spermatogenesis requires a type of testis-specific 20S core proteasome. Nat Commun 10, 3387 (2019).

15. T. Menneteau, et al., Mass spectrometry-based absolute quantification of 20S proteasome status for controlled ex-vivo expansion of Human Adipose-derived Mesenchymal Stromal/Stem Cells. Molecular & cellular proteomics: MCP, mcp.RA118.000958 (2019).

16. A. Kniepert, M. Groettrup, The unique functions of tissue-specific proteasomes (2014).

17. M.-S. Kim, et al., A draft map of the human proteome. Nature 509, 575–581 (2014).

18. L. Gómez-H, et al., The PSMA8 subunit of the spermatoproteasome is essential for proper meiotic exit and mouse fertility. PLoS Genet 15, e1008316 (2019).

19. Z.-H. Zhang, et al., Proteasome subunit α4s is essential for formation of spermatoproteasomes and histone degradation during meiotic DNA repair in spermatocytes. J. Biol. Chem., jbc.RA120.016485 (2020).

20. J. S. Ahuja, et al., Control of meiotic pairing and recombination by chromosomally tethered 26S proteasome. Science 355, 408–411 (2017).

21. F. Bassermann, R. Eichner, M. Pagano, The ubiquitin proteasome system - implications for cell cycle control and the targeted treatment of cancer. Biochim Biophys Acta 1843, 150–162 (2014).

22. X. Gao, et al., The REGγ-Proteasome Regulates Spermatogenesis Partially by P53-PLZF Signaling. Stem Cell Reports 13, 559–571 (2019).

23. B. Fabre, et al., Deciphering preferential interactions within supramolecular protein complexes: the proteasome case. Molecular systems biology 11, 771 (2015).

24. D. Sbardella, et al., The insulin-degrading enzyme is an allosteric modulator of the 20S proteasome and a potential competitor of the 19S. Cellular and Molecular Life Sciences, 1–16 (2018).

25. J. Lesne, M.-P. Bousquet, J. Marcoux, M. Locard-Paulet, Top-Down and Intact Protein Mass Spectrometry Data Visualization for Proteoform Analysis Using VisioProt-MS. Bioinform Biol Insights 13, 1177932219868223 (2019).

26. B. Guillaume, et al., Two abundant proteasome subtypes that uniquely process some antigens presented by HLA class I molecules. Proceedings of the National Academy of Sciences 107, 18599–18604 (2010).

27. B. Fabre, et al., Label-Free Quantitative Proteomics Reveals the Dynamics of Proteasome Complexes Composition and Stoichiometry in a Wide Range of Human Cell Lines. Journal of Proteome Research 13, 3027–3037 (2014).

28. M. Locard-Paulet, et al., VisioProt-MS: interactive 2D maps from intact protein mass spectrometry. Bioinformatics 35, 679–681 (2019).

29. J. D. Thompson, C. Schaeffer-Reiss, M. Ueffing, Functional proteomics. Preface. Methods Mol Biol 484, v–vii (2008).

30. S. Vimer, et al., Comparative Structural Analysis of 20S Proteasome Ortholog Protein Complexes by Native Mass Spectrometry. ACS Cent Sci 6, 573–588 (2020).

31. H. Uechi, J. Hamazaki, S. Murata, Characterization of the testis-specific proteasome subunit ??4s in mammals. Journal of Biological Chemistry 289, 12365–12374 (2014).

32. M.-P. Bousquet-Dubouch, et al., Affinity Purification Strategy to Capture Human Endogenous Proteasome Complexes Diversity and to Identify Proteasome-interacting Proteins. Molecular & Cellular Proteomics 8, 1150–1164 (2009).

33. T. A. Griffin, et al., Immunoproteasome assembly: cooperative incorporation of interferon gamma (IFN-gamma)-inducible subunits. The Journal of Experimental Medicine 187, 97–104 (1998).

34. D. J. Kingsbury, T. A. Griffin, R. A. Colbert, Novel Propeptide Function in 20 S Proteasome Assembly Influences Subunit Composition. Journal of Biological Chemistry 275, 24156–24162 (2000).

35. S. Heink, D. Ludwig, P.-M. Kloetzel, E. Krüger, IFN-γ-induced immune adaptation of the proteasome system is an accelerated and transient response. PNAS 102, 9241–9246 (2005).

36. H. K. Johnston-Carey, L. C. D. Pomatto, K. J. A. Davies, The Immunoproteasome in oxidative stress, aging, and disease. Critical Reviews in Biochemistry and Molecular Biology 51, 268–281 (2016).

37. X. Li, D. Thompson, B. Kumar, G. N. DeMartino, Molecular and Cellular Roles of PI31 (PSMF1) Protein in Regulation of Proteasome Function*. Journal of Biological Chemistry 289, 17392–17405 (2014).

38. R. Kirk, et al., Structure of a conserved dimerization domain within the F-box protein Fbxo7 and the PI31 proteasome inhibitor. J Biol Chem 283, 22325–22335 (2008).

39. C. C. Rathje, et al., A Conserved Requirement for Fbxo7 During Male Germ Cell Cytoplasmic Remodeling. Frontiers in Physiology 10, 1278 (2019).

40. M. Bader, et al., A conserved F box regulatory complex controls proteasome activity in Drosophila. Cell 145, 371–382 (2011).

41. L. Huang, K. Haratake, H. Miyahara, T. Chiba, Proteasome activators, PA28γ and PA200, play indispensable roles in male fertility. Scientific Reports 6 (2016).

42. X. Wang, L. Huang, Identifying dynamic interactors of protein complexes by quantitative mass spectrometry. Mol Cell Proteomics 7, 46–57 (2008).

43. B. Fabre, et al., Subcellular Distribution and Dynamics of Active Proteasome Complexes Unraveled by a Workflow Combining in Vivo Complex Cross-Linking and Quantitative Proteomics. Molecular & Cellular Proteomics 12, 687–699 (2013).

44. B. Khor, et al., Proteasome activator PA200 is required for normal spermatogenesis. Mol Cell Biol 26, 2999–3007 (2006).

45. B. M. Stadtmueller, C. P. Hill, Proteasome Activators. Molecular Cell 41, 8–19 (2011).

46. J. Lesne, et al., Conformational maps of human 20S proteasomes reveal PA28- and immuno-dependent inter-ring crosstalks. Nature Communications 11, 6140 (2020).

47. H. B. D. P. Rao, et al., A SUMO-ubiquitin relay recruits proteasomes to chromosome axes to regulate meiotic recombination. Science 355, 403–407 (2017).

48. R. Bose, G. Manku, M. Culty, S. S. Wing, Ubiquitin-proteasome system in spermatogenesis. Adv Exp Med Biol 759, 181–213 (2014).

49. A. Minis, et al., The proteasome regulator PI31 is required for protein homeostasis, synapse maintenance, and neuronal survival in mice. PNAS 116, 24639–24650 (2019).

50. P. F. Cho-Park, H. Steller, Proteasome regulation by ADP-ribosylation. Cell 153, 614–627 (2013).

51. K. Liu, et al., PI31 Is an Adaptor Protein for Proteasome Transport in Axons and Required for Synaptic Development. Developmental Cell 50, 509–524.e10 (2019).

52. E. K. Meziane, S. J. Randle, D. E. Nelson, M. Lomonosov, H. Laman, Knockdown of Fbxo7 reveals its regulatory role in proliferation and differentiation of haematopoietic precursor cells. J Cell Sci 124, 2175–2186 (2011).

53. J. Blickwedehl, et al., Role for proteasome activator PA200 and postglutamyl proteasome activity in genomic stability. Proceedings of the National Academy of Sciences 105, 16165–16170 (2008).

54. Y. Perez-Riverol, et al., The PRIDE database and related tools and resources in 2019: improving support for quantification data. Nucleic Acids Res 47, D442–D450 (2019).

