## Supplementary material for "Proteasome complexes experience profound structural and functional rearrangements throughout mammalian spermatogenesis": Fig. S1

### Supporting Information

#### SI Experimental Procedures

Antibody development

Two peptides were synthesized, C+ILNPEEIEKYVAEIEKEKEENEKKKQKKAS - corresponding to the α4-specific C-terminal sequence (P1 peptide) and C+MFSAKEIELQVNEIEKEKEEAEKKKSKKTT - corresponding to the α4s-specific C-terminal sequence (P2 peptide). Two New Zealand White rabbits, 20 to, 25 weeks old, were immunized with a KLH-P2 conjugate (Day 0), by subcutaneous injection containing incomplete Freund's adjuvant. Immunisations were repeated on Day 7, Day 14 and Day 35. The rabbits were exsanguinated on Day 42 after the immunisation and the antibodies were affinity-purified from the blood serum using the immobilized P1 peptide, followed by depletion step using immobilized P2 peptide, in order to remove the α4-reactive antibodies. The obtained anti-α4s polyclonal antibodies exhibited an EC50 of 0.005 µg/mL against the α4s epitope and of 2.12 µg/mL against its α4 counterpart (**Fig.S4B**). Moreover, the specificity of the newly-produced anti-α4s antibody was verified by western-blot assay (**Fig.S4C**).

Preparation of separated germ cells from rats

Spermatogonia (SPG) were isolated from testes obtained from thirty 8-day-old Sprague Dawley (SD) rat testes with a purity of greater than 90% as previously described (1). Seminiferous epithelial cells were dispersed by enzyme treatment and separated. Briefly, the cells were separated by sedimentation velocity at unit gravity at 4 °C, using a 2–4% BSA gradient in Ham'F12/DMEM in an SP-120 chamber (STAPUT). After 2.5 h of sedimentation, 35 fractions were collected. Cell fractions 16 to 21 were pooled, washed 3 times with Phosphate Buffered Saline pH 7.4 and cells were immediately subjected to in vivo cross-linking to stabilize proteasome interactors, as previously described (2). Cross-linked cells were stored at -80°C.

Pachytene spermatocytes (SPC) and early spermatids (SPT) were isolated from testes obtained from eight 90-day-old SD rat and prepared by centrifugal elutriation with a purity greater than 90% according to a previously described method (3), except that enzymatic dissociation of cells was replaced by mechanical dispersion. Upon isolation, cells were washed 3 times with Phosphate Buffered Saline pH 7.4 and cells were immediately subjected to in vivo cross-linking as previously described (2). Cross-linked cells were stored at -80°C.

Sertoli cells (SER) were isolated from testes obtained from eight 20-day-old SD rats according to previously described methods (4, 5). The cell suspensions were seeded at a density of approximately 1x10^6 cells/mL in 75 cm2 tissue culture flasks (NUNC, Copenhagen, Den- mark). The cells were then incubated at 32°C in a humidified atmosphere of 5% $CO_2$ and 95% air in Ham's F12/ DMEM supplemented with insulin (10 mg/mL), Human transferrin (5 mg/mL) and gentamycin (10 mg/mL), all from Life Technologies (Eragny, France). Culture medium was changed daily until the end of the experiment. On the second day of culture, the cells were exposed to a hypotonic treatment in order to eliminate the contaminating germ cells. The degree of purity of the isolated Sertoli cells was > 98%. On day 5 of culture, cells were recovered, washed 3 times with Phosphate Buffered Saline pH 7.4 and were immediately subjected to in vivo cross-linking as previously described (2). Cross-linked cells were stored at -80°C.

Proteasome Purification

The proteasome complexes were immunopurified from tissues or cell lysates using the MCP21 antibody as described before (6), the anti-α4s or anti-PA200 antibodies as fully described in SI Experimental Procedures. Protein extract were trypsin digested using S-trap micro cartridges (Protifi, Farmingdale NY USA)

Eight mg of MCP21 antibody was grafted to one gram of CNBr-activated Sepharose beads. Frozen bovine testis (-80 °C) from an adult individual were broken into pieces using a hammer and chisel. Lysis and coIP were then done at 4 °C. Ten grams of tissue were diced and placed into a blender with the lysis buffer (20 mM Tris HCl, 0.25% TritonX-100, 100 mM NaCl, 10 mM ATP, 5 mM $MgCl_2$, 1 tablet of inhibitors/50 mL (cOmplete™ ULTRA Tablets Mini EDTA-free, Merck, Darmstadt, Germany), pH 7.6), at 1.6 weight to volume ratio. The lysate was passed through a Dounce homogenizer and then sonicated in a VibraCell (Bioblock Scientific, France) (power = 2, 60% active cycle, 10 cycles, 30 sec per cycle). The lysates were centrifuged for 30 min at 4 °C, 4500 g and the greasy supernatant was aspirated from the top of the lysate. The solid pellet was also discarded while the remaining liquid was again centrifuged at 4 °C and 16000 g, for 30 min. The supernatant was filtered through a 0.22 µm membrane. The filtered lysate was then mixed with the Sepharose grafted with antibodies and incubated at 4°C overnight. The next day, the Sepharose was washed with 50 mL of Equilibration buffer (20 mM Tris HCl, 1 mM EDTA, 10% glycerol, 100 mM NaCl, pH 7.6) and eluted with Equilibration buffer supplemented with 3 M NaCl. The eluate was concentrated on Amicon filters of 100 kDa cutoff (Millipore) and separated using gel filtration chromatography. The chromatography was performed on a Superose 6 10/300 GL column, using TSDG buffer (10 mM Tris-HCl pH 7.0, 1 M KCl, 10 mM NaCl, 5.5 mM $MgCl_2$, 0.1 mM EDTA, 1 mM DTT, 10% Glycerol). The fractions were assayed by SDS-PAGE and 20S proteasome fractions were pooled together.

Proteasome immunopurification from rat germ cells and sample preparation for LC-MS

Immunopurification of proteasome complexes from rat germs cells was done using the MCP21 clone of the anti-α2 antibody and sample preparation was done by in-gel digestion, as described in (7). For this protocol, pools of cells from different rats, 8-10x10^6 SER, 20-25x10^6 of SPGs, 8x10^6 of SPCs and 20x10^6 of SPTs per replicate were used. Three independent experiments were run for each type of cell.

Co-immunopurification for interactomics from bovine testis and sample preparation for LC-MS/MS

Four biological replicates of frozen bovine testes (-80 °C) from adult individuals were broken into pieces using a hammer and chisel. Lysis and coIP were then done at 4 °C. Ten grams of tissue were diced and placed into a blender with the lysis buffer (20 mM Tris HCl, 0.25% TritonX-100, 100 mM NaCl, 10 mM ATP, 5 mM MgCl2, 1 tablet of protease inhibitors/50 mL (cOmplete™ ULTRA Tablets Mini EDTA-free, Merck, Darmstadt, Germany), pH 7.6), 1.6 weight to volume ratio. The lysate was passed through a Dounce homogenizer and then sonicated in a VibraCell (Bioblock Scientific, France) (power=2, 60% active cycle, 10 cycles, 30 sec per cycle). The lysates were centrifuged for 30 min at 4 °C, 4500 g and the greasy supernatant was aspirated from the top of the lysate. The solid pellet was also discarded while the remaining liquid was again centrifuged at 4°C and 16,000 g for 30 min. The supernatant was filtered through a 0.22 µm membrane. The co-immunopurification of the proteasome complexes was done using ProteinG coupled to the magnetic beads (MagBeads, Cat. No. L00274, GenScript USA Inc. Piscataway, NJ USA). The protocol proposed by the supplier was followed with the following modifications. The amount of beads used per sample was 100 µL of suspension. The amount of antibodies used per replicate was 100 µg of anti-α4s, 50 µg of anti-α2, 10 µg of anti-PA200 and 50 µg of OX8 control antibody. One mL of lysate of each replicate was incubated with ProteinG-MagBeads, coupled with different antibodies, overnight at 4 °C on a rocker. Samples were then washed four times with Equilibration buffer (20 mM Tris HCl, 1 mM EDTA, 10% glycerol, 100 mM NaCl, 2 mM ATP, 5 mM MgCl2, pH 7.6), then eluted with 25 µL of S-trap Elution Buffer (5% SDS, 50 mM TEAB, pH 7.55). In case of anti-α4s, proteasome complexes were eluted via competitive elution method, for improved signal/noise ratio. Namely, the peptide used for immunization of the animals and production of antibody was used to elute the antibody from the MagBeads. Peptide was diluted in the Equilibration Buffer to 160 µg/mL and 100 µL of the peptide solution were added per replicate. The beads were incubated for 1h at room temperature, while mixing. The supernatant was recovered and mixed with SDS solution to final 5% SDS. Control IP was also done with the OX8 control antibody and the peptide elution procedure.

Preparation for LC-MS analysis was done using S-trap micro cartridges (Protifi, Farmingdale NY USA), by default protocol of the manufacturer. Around 2 µg of peptides per sample per run were used for LC-MSMS analysis.

Bottom-up LC-MSMS

Analysis was performed on an Orbitrap Fusion instrument, with nano ESI source, coupled to the Ultimate 3000 chromatography system using 90 min runs. Peptides were loaded onto a precolumn and washed (97.95% water, 2% ACN,0.05% trifluoroacetic acid). Upon loading onto the column (made in-house, 75µm ID, 50cm column, packed with Reprosil C18, 3 µm phase, Cluzeau), the elution gradient is initiated at 300 µL/min, by 100% phase A (0.2% formic acid) for 3min, then a slope 10-30% phase B (80% acetonitrile, 0.2%

formic acid) for 50min, followed by 30-45% B gradient for 10min, to finish with 45-80% B during 1min. The 80% B was maintained for 10 to wash the column, followed by re-equilibration with the 100% phase A during 15min. The nano- ion source was at +1900 V, with ion transfer tube temperature at 275 °C. MS analysis was done in the Orbitrap at 60k resolution, scan range between 350 and 1400 m/z, with RF lens value set to 60%, AGC target set to 4e5, maximum injection time to 50 ms, with 1 microscan performed in profile mode. MSMS was done in data-dependant mode, in the Orbitrap, at resolution of 15k (first mass at 100 m/z). Monoisotopic peak determination was set for peptides, intensity threshold of 2.5e4 was used and only charge states from 2 to 6 were used as precursors. Dynamic exclusion was used with the following parameters: exclude the ion after “1” times, during 30 sec, including isotopes, with mass tolerance, high and low, 10 ppm each. Ion isolation was done in the quadrupole, with an isolation window of 1.7 m/z. The activation was done by HCD at 28% collision energy. AGC was set to 5e4, ions were injected for all available parallelizable time, with maximum injection time of 22 ms. A total of 1 microscans per cycle was used and recorded in centroid mode.

Bottom-Up data treatment

Identification and quantitation parameters. The identification was done with Mascot search engine, using following search parameters: Peptide mass tolerance : 10 ppm; Peptide charge : 2+ and 3+; Cleavage enzyme: Trypsin/P; Max. missed cleavages : 2; Decoy : true; Mass : Monoisotopic; Fixed modifications : Carbamidomethyl (C); Variable modifications : Acetyl (Protein N-term),Oxidation (M); MS/MS IONS SEARCH : true; Error tolerant : false; MS/MS tolerance : 20 mmu; Data format : Mascot generic; Quantitation : None; Instrument : ESI FTMS HCD. The UniProt entries (SwissProt & TrEMBL) for *Bos Taurus* were used as a protein entry database, including isoforms. In the case of rat germ cells, the search was performed with *Rattus norvegicus* Uniprot database, including isoforms.

The result files were then imported into Proline – software for Bottom-Up proteomics (8), using following parameters: Peaklist software: Proline; Generate intermediate mzIdentML file: false; Instrument configuration: ORBITRAP FUSION (A1=FTMS F=HCD A2=FTMS); Decoy strategy: Concatenated Decoy Database; Protein match decoy rule: SearchGUI REVERSED; Ion score cutoff: None; Subset threshold: None; Replace peptide matches scores by PEP values (Posterior Error Probabilities): false; Automatically associate raw files to identification results (DAT, OMX): true. Validation of the peptide IDs in Proline was done using following parameters: PSM filters: Pretty Rank: 1.0, Peptide Seq Length: 7.0; PSM validation: expecting FDR of 1.0% on Mascot Adjusted E-Value; Validation of the protein sets was done using the following parameters: Min. specific peptide count: 1; Enable protein FDR (false discovery rate) filter: true; Min protein set FDR (%): 1.0; Peptide set scoring method: Mascot Modified MudPIT; Dataset merging mode: After validation (merge identification summaries). Label-free quantitation was done using following parameters: Quantitation type: label_free; Quantitation method: Label free based on the extraction of feature abundance; Extraction m/z tolerance: 5 ppm; Mapping m/z tolerance:

5 ppm; Min. peakel duration: 15 sec; Alignment computation method: (Iterative; Max number of iterations: 3; m/z tolerance: 5 ppm; Time tolerance: 5 sec); Alignment smoothing: (Method: Landmark Range; Number of landmarks: 50; Sliding window overlap: 50); Features cross assignment: (Mapping m/z tolerance: 5 ppm; Time tolerance: 60 sec; Normalization method: None; Master feature filter type: Intensity; Intensity threshold: 0). Normalization and statistics were done using the following parameters: Global options: (Apply profile clustering: false (method: QUANT_PROFILE); Use only specific peptides: true; Discard miscleaved peptides: false (method: KEEP_MOST_ABUNDANT_FORM); Discard oxidized peptides: false (method: KEEP_MOST_ABUNDANT_FORM); Abundance summarizer method: LFQ based on QUANT_PEPTIDE_IONS); Protein options: (Apply normalization: true; Apply missing values inference: false; Apply variance correction: false; Apply T-test: true (p-value: 0.01); Apply Z-test: true (p-value: 0.4)); Peptide options: Apply normalization: true; Apply missing values inference: false; Apply variance correction: false; Apply T-test: true (p-value: 0.01); Apply Z-test: true (p-value: 0.4). The p-value for a non-paired, one-sided T-test and fold change were calculated between the experimental IP groups and the control IP groups (IP with the OX8 antibody). Proteins enriched over 2-fold with a p-value below 0.05 were considered significantly enriched.

Comparison between the anti- α2 and anti- α4s was done in the following way:

(1) Datasheet from the Proline export file, containing all the protein set entries with their relative abundance, called “Protein sets” was copied for treatment and the empty fields were replaced with zeros.

(2) Logical tests for every protein entry were performed, to determine if it was detected in at least 4 out of 8 analysis runs of at least one group of samples (4 samples times 2 technical replicates), then that it was determined by specific peptides, then filtered only the entries that satisfy the said tests. The logic tests were done on raw intensities, not the normalized ones.

(3) The technical replicates were averaged and the fifth percentile for every group was calculated.

(4) Number of missing values for each IP sample were calculated and the missing values were replaced by the fifth percentile for protein entries with two or more missing values.

(5) The iBAQ values were calculated by dividing the intensities with the number of observable peptides for each entry.

(6) Mean proteasome abundance was calculated as average of iBAQs for non-catalytic subunits of the proteasome, excluding the α4 and α4s.

(7) The iBAQ values were normalized in all the groups with mean proteasome value for each group.

(8) The one-sided paired T-test and fold change were calculated between the anti-α2 and anti-α4s groups. The comparison was done only for proteins significantly enriched between the experimental and control groups (see the end of previous paragraph).

<u>Estimation of the regulator stoichiometry of the α4- and α4s-containing proteasome :</u>

The composition of these two complexes can be estimated for known interactors based on their intensities in the two CoIPs that have a different ratio of enrichment of α4 and α4s, which is the case in our experiments (anti-α2 and anti-α4s CoIP). Firstly, we need to know the ratio of α4 versus α4s in both CoIPs. The intensity of an interactor of a particular complex (α4- or α4s-containing proteasome) can be expressed via the following formulas:

$x=(A(\alpha2)*d - A(\alpha4s)*b)/(a*d - b*c)$

$y=(A(\alpha2)*c - A(\alpha4s)*a)/(b*c - a*d)$

if starting from statements:

$A(\alpha2)=a*x+b*y$ and $A(\alpha4s)=c*x+d*y$

for a,b,c,d being the percentages in 0 to 1 format (e.g. 0.15 for 15%), where x is the intensity of a protein P bound to α4-containing P20S and y is the intensity of a protein P bound to α4s-containing P20S. A(α2) is the experimentally measured intensity of the protein P in the anti-α2 CoIP, while A(α4s) is the experimentally measured intensity of the protein P in the anti-α4s CoIP.

Additionally:

since $a=1-b$ and $c=1-d$

then:

$x=(b*A(\alpha4s) - d*A(\alpha2))/(b-d)$

$y=((1-d)*A(\alpha2) + (b-1)*A(\alpha4s))/(b-d)$

(conditions: $b \neq d$, $bd \neq d$)

The estimations are easily derived when the two formulae are solved together, once the intensity values of a protein P for each CoIP are plugged in, together with the α4s percentage in each CoIP (b and d factors, 42% and 96% for anti-α2 and anti- α4s IP, respectively).

#### Experimental Design and Statistical Rationale

Label-free MS Analyses: All statistical analyses were performed on at least three independent biological replicates. For each biological replicate, results from at least two injection replicates were averaged. The analyses on bovine testes were done in four biological replicates. Probability values (p) were determined by one-way paired Student's test for groups of equal variance. Differences were statistically significant at confidence levels of 95% (*), 99% (**), or 99.9% (***). The Pull-Down assays were done in three technical replicates, with p values determined by two-sided, non-paired Student's test for groups of equal variance.

The "rat germ cells" dataset contains MS results from the analysis of the IPs and lysates of 4 cell types (Sertoli cells, spermatogonia, spermatocytes, spermatids) in biological triplicates (pools of cells from different individuals for each group), analysed in three injection replicates, for a total of 72 raw files. The "bovine testis CoIP" dataset contains MS results for 4 types of IPs (anti-PA200, anti-α2, anti-α4s, and OX-8 control) from four biological replicates, two technical injections per biological replicate, and lysates for each biological sample, for a total of 40 .raw files. The quantitative comparison of proteins was done based on normalized protein intensities from the "Protein groups" sheet in the Proline output file.

#### Top-down LC-MSMS

Nano-LC-MS analyses of commercial or immunopurified 20S were performed on a nanoRS UHPLC system (Dionex) coupled to an LTQ-Orbitrap Fusion Tribrid mass spectrometer (Thermo Fisher Scientific), as previously described (9). Briefly, a total of 5 µL of sample at 0.5 µM was loaded onto a reverse-phase C4-precolumn (300 µm i.d. × 5 mm; Thermo Fisher Scientific) at 20 µL/min in 2% acetonitrile (ACN) and 0.2% formic acid (FA). After 5 minutes of desalting, the precolumn was switched online with an analytical C4 nanocolumn (75 µm i.d. × 15 cm; in-house packed with C4 Reprosil) equilibrated in 95% solvent A (5% ACN, 0.2% FA) and 5% solvent B (0.2% FA in ACN). Proteins were eluted using a binary gradient ranging from 5% to 40% (5 min) and then 40% to 99% (33 min) of solvent B at a flow rate of 300 nL/min. The Fusion Tribrid (Thermo Fisher Scientific) was operated in single MS acquisition mode with the Xcalibur software (Thermo Fisher Scientific). The spray voltage was set to 1900 V, the ion transfer tube temperature to 350°C, the RF lens to 60%, and no in-source dissociation was applied. The MS scans were acquired in the 700 to 2000 m/z range with the resolution set to 15 000 and using 10 µscans for averaging. LC-MS .raw files were automatically deconvoluted with the rolling window deconvolution software RoWinPro (10) and the proteoform footprints were visualized with VisioProt-MS v2.0 (11).

For MSMS acquisition, 3 second cycles were used, with the following MS parameters: MS scan range between 1000 and 2000 m/z in Orbitrap at 7,500 resolution, 10 µscans for averaging. MSMS activation (HCD) was done with 25% activation energy, with spectra acquisition in the Orbitrap at 60,000 resolution. The AGC target was set to 1e6. Dynamic exclusion for 600 s was used within 60 s to prevent repetitive selection of the same precursor (selection tolerance: ±10ppm) and improve the number of identified proteins. The following source settings were used: Spray voltage 1900V, RF lens 60%, no in-source fragmentation was used. The MSMS analysis of the degraded 20S proteasome used the same parameters, except for the activation method, whereby EThcD was used instead of HCD: ETD activation time of 5ms, ETD reagent target intensity set at 7e5, maximum ETD reagent injection time set at 200ms, with supplemental EThcD activation at 25% energy. The spectra identification was done in Proteome Discoverer (Thermo Fisher Scientific) v.2.3, using an Absolute Mass Search with the following parameters: precursor mass tolerance - 1000 Da, fragment mass tolerance - 15ppm. Mutations and truncations of the proteoforms were then manually curated. The search database was generated from the ensemble of the available Swiss-Prot and TrEMBL entries for bovin 20S proteasome, bovin proteasome regulators (19S, PA28 subunits, and PA200) and their isoforms.

HDX-MS and MS simulations

HDX-MS experiments were performed on a Synapt-G2Si (Waters Scientific, Manchester, UK) coupled to a Twin HTS PAL dispensing and labelling robot (Trajan Scientific, Milton Keynes, UK) via a NanoAcquity system with HDX technology (Waters, Manchester, UK). Data were collected with MassLynX v.4.1 (Waters Scientific (Manchester, UK) and the robot was controlled via HDX Director v.1.0.3.9 (LEAP Technologies, Carborro, NC, USA). Each step was optimized to minimize sample loss and work with such a heterogeneous sample, as described previously (12).

Briefly, 5.7 µL of bovin 20S proteasome at 2.3 µM were aspirated and 5.2 µL were diluted in 98.8 µL of protonated (peptide mapping) or deuterated buffer (20 mM Tris pH/pD 7.4, 1 mM EDTA, 1 mM DTT) and incubated at 20 °C for 0, 0.5, 1, 5, 10, and 30 min. The final D2O percentage was 95% in the labelled samples. 99 µL were then transferred to vials containing 11 µL of precooled quenching solution (500 mM glycine at pH 2.3). After 30 s. of quenching, 105 µL were injected into a 100 µL loop. Proteins were digested online with a 2.1 × 30 mm Poros Immobilized Pepsin column (Life Technologies/Applied Biosystems, Carlsbad, CA, USA). The temperature of the digestion room was set at 15 °C.
Peptides were desalted for 3 min on a C18 pre-column (Acquity UPLC BEH 1.7 µm, VANGUARD) and separated on a C18 column (Acquity UPLC BEH 1.7 µm, 1.0 × 100 mm) by the following gradient: 5–35% buffer B (100% acetonitrile, 0.2% formic acid) for 12 min,

35–40% for 1 min, 40–95% for 1 min, 2 min at 95% followed by 2 cycles of 5–95% for 2 min and a final equilibration at 5% buffer A (5% acetonitrile, 0.2% formic acid) for 2 min. The total runtime was 25 min. The temperature of the chromatographic module was set at 4 °C. Experiments were run in triplicates and the protonated buffer was injected between each triplicate to wash the column and avoid cross-over contamination.
The acquisitions were performed in positive and resolution mode in the m/z range 50–2000 Th. The sample cone and capillary voltages were set at 30 and 3 kV, respectively. The analysis cycles for nondeuterated samples alternated between a 0.3 s low energy scan (Trap and Transfer collision energies set to 4 V and 2 V, respectively), a 0.3 sec high energy scan (Ramp Trap and Transfer collision energies set to 18 to 40 V and 2 to 2 V, respectively) and a 0.3 sec lockspray scan (0.1 µM [Glu1]-Fibrinopeptide in 50% acetonitrile, 50% water and 0.2% formic acid infused at 10 µL/min). The lockspray trap collision energy was set at 32 V and a GFP scan of 0.3 sec is acquired every min. Deuterated samples were acquired only with the low energy and lockspray functions.
Peptide identification was performed with ProteinLynx Global SERVER v.3.0.2 (PLGS, Waters, Manchester, UK) based on the MSE data acquired on the nondeuterated samples. The MSMS spectra were searched against a home-made database containing sequences of the 18 c20S, i20S and s20S subunits, all the 19S subunits, as well as the entries for PA28αβ, PA28γ and PA200 regulators, as well as pepsin from Sus scrofa. Peptides were filtered in DynamX v.3.0 from Waters Scientific (Manchester, UK) with the following parameters: peptides identified in at least two replicates, 0.2 fragments per amino-acid, intensity threshold 1000. The RDUs were not corrected for back exchange. These data were then used for structural data mining with HDX-Viewer (13). Molecular representations of **Fig.5** were generated in UCSF ChimeraX v.0.9 (14) and Visual Molecular Dynamics (15), respectively.

Molecular Dynamics simulations and analysis

For molecular dynamics (MD simulations), we used the α4 model to perform an homology model of the bovine α4s subunit using the modeler program (16) (83.2 % sequence identity between α4 and α4s). For each structure, we then performed 1 µs of molecular dynamics (MD) simulations using GROMACS 2020 (17) in combination with AMBER14SB force field (18). We first embedded each model in a waterbox using TIP3P water model (19). Cl ions were added to neutralize the system. Energy minimization was performed using the steepest descent algorithm and each system was equilibrated with a constant temperature (canonical ensemble, NVT, 300 K) ensemble for 100 picoseconds (ps), followed by a 100 ps equilibration at constant pressure (isothermal-isobaric, NPT, 1 bar). For equilibration steps, the protein backbone was kept constrained. For equilibration and production runs, we applied the velocity-rescaling thermostat (20) on protein and solvent, coupled with the Parrinello–Rahman barostat (21), with a time constant of 2 ps and compressibility of 4.5x10-5 bar-1. Long-range electrostatics were modeled using the

Particle-Mesh Ewald method (22, 23). All bonds were treated using the LINCS algorithm (24). For both α4 and α4s models, we performed 1 µs of simulation at 300 K and ambient pressure of 1 bar with an integration time step of 2 femto-seconds.

We then analyzed the dynamics of the two last α-helices. We analyzed the creation of hydrogen bonds between water molecules and helices NH groups. To do so, we used VMD (15) Hbonds plugin (https://www.ks.uiuc.edu/Research/vmd/plugins/hbonds/) to count hydrogen bonds formed every 100 ps of the trajectory using a donor-acceptor distance of 3.5 Å and a cutoff angle of 30°. We then calculated the Solvent Accessible Surface Area (SASA) using VMD functions (https://www.ks.uiuc.edu/Research/vmd/vmd-1.8.3/ug/node121.html). To be comparable with HDX-MS data, only the NH groups of residues 180 to 189 and 225 to 241 were used for these two analyses. We calculated hydrogen bonds formed between the two helices (respectively residues 185 to 198 and 224 to 241) as well as the inter-helix angle θ every 100 ps. We used VMD Hbonds plugin and Tcl functions to calculate respectively interhelix hydrogen bonds and interhelix angle.

Activity assays

The activity assay was performed in 96-well black plates (Greiner Bio- One, UK). For assays on purified proteins, 11 µL of purified 20S proteasome sample at 0.57 µM (either c20S or s20S) was mixed with 17.75 µL of recombinant PA200 regulator at 2.84 µM and left to interact for 30 min at 37°C. After they were allowed to interact, the complexes were diluted with activity buffer (20 mM Tris HCl, 5 mM MgCl2, 2 mM ATP, 0.5 mM DTT) to a final volume of 425 µL. As a control, 11 µL of purified 20S proteasome sample at 0.57 µM (either c20S or s20S) was also diluted to a final volume of 425 µL. All the mixes were distributed into wells in aliquots of 50 µL (except for the assays of LLVY, where 25 µL of sample + 25 µL of Activity buffer were used) to which 50 µL of Activity buffer was added, supplemented with 100 µM fluorogenic peptide substrate: Suc-LLVY-AMC, Boc-LRR-AMC or z-LLE-AMC, to probe for chymotrypsin-, trypsin- or caspase-like activities, respectively. The kinetic assays were performed at 37 °C in a FLX-800 spectrofluorometer (BIOTEK) over 60 min with one reading every 5 min, at 360 nm for excitation and 460 nm for emission. The slope of the kinetic assay (increase in fluorescence intensity over time) and the quantity of proteasome were used to determine the proteasome’s specific activity.

Pull-Down Assays

20S proteasomes were purified from bovine testes as already described above. It is established that the 20S pool contains ~50% s20S and ~50% α4-containing 20S. Purified recombinant human 19S complex was bought from R&D Systems (Minneapolis, USA), recombinant, FLAG-tagged human PI31 protein was bought from OriGene (Rockville,

USA) and PA200 protein was prepared as described in (Toste Rêgo and da Fonseca, 2019).
The PI31 and 20S pool were mixed at a 10 to 1 ratio in Equilibration buffer 1 (20 mM Tris HCl, 1 mM EDTA, 10% glycerol, 100 mM NaCl, 2 mM ATP, 5 mM MgCl2, pH 7.6) in a total volume of 100µL, at final concentrations of 280nM 20S and 2.8µM PI31. The mix was incubated for 3,5h at 4°C with either the anti-FLAG resin or the proteinG MagBeads containing 100µg of anti-α2 antibody. The resin or the beads were washed with 5mL Equilibration buffer. The elution was done with 2x volume of the slurry of Glycine buffer (Glycine pH3,5 0,1 M) and the samples were prepared for LC-MS analysis on mini-S-trap cartridges (Protifi, Farmingdale NY USA) as recommended by the supplier. For FLAG-tag IP, 280 µL of 50% M2 anti-FLAG affinity resin slurry was used (Merck, Darmstadt, Germany). For anti-α2 IP, 100 µL of 25% MagBeads slurry were used (GenScript, Piscataway, NJ USA).
The PA200 of 19S were separately mixed with 20S pool in a 1:2 ratio (10 µg of 19S with 20 µg of 20S proteasome and 2.94 µg PA200 with 20 µg 20S proteasome) in the Equilibration buffer 2 (50 mM Tris pH 7.6, 1 mM MgCl2, 200 µM ATP, 0.5 M DTT) in a total volume of 24 µL. Both mixes were incubated for 30 min at 37 °C, then split in two halves. One half was mixed with the experimental group antibody (10 µg of anti-PSMC2 for 19S-20S mix or anti-PA200 for PA200-20S mix) or control group antibody (10 µg of anti-α2 antibody) and incubated for 30 min at 37°C. Each of the samples was then mixed with ProteinG MagBeads (50 µL 25% slurry per sample) and incubated overnight at 4 °C. The elution of complexes was done with 50 µL of S-Trap buffer and the samples were prepared for LC-MS analysis on mini-S-trap cartridges (Protifi, Farmingdale NY USA) as recommended by the supplier.

**Additional References:**


1. E. Com, B. Evrard, P. Roepstorff, F. Aubry, C. Pineau, New Insights into the Rat Spermatogonial Proteome. *Molecular & Cellular Proteomics* **2**, 248–261 (2003).
2. B. Fabre, *et al.*, Subcellular Distribution and Dynamics of Active Proteasome Complexes Unraveled by a Workflow Combining in Vivo Complex Cross-Linking and Quantitative Proteomics. *Molecular & Cellular Proteomics* **12**, 687–699 (2013).
3. C. Pineau, V. Syed, C. W. Bardin, B. Jégou, C. Y. Cheng, Germ cell-conditioned medium contains multiple factors that modulate the secretion of testins, clusterin, and transferrin by Sertoli cells. *J Androl* **14**, 87–98 (1993).
4. M. K. Skinner, I. B. Fritz, Testicular peritubular cells secrete a protein under androgen control that modulates Sertoli cell functions. *Proceedings of the National Academy of Sciences of the United States of America* **82**, 114–118 (1985).
5. A. M. W. Toebosch, D. M. Robertson, I. A. Klaij, F. H. De Jong, J. A. Grootegoed, Effects of FSH and testosterone on highly purified rat Sertoli cells: inhibin ?-subunit mRNA expression and inhibin secretion are enhanced by FSH but not testosterone. *Journal of Endocrinology* **122**, 757–762 (1989).
6. B. Fabre, *et al.*, Label-Free Quantitative Proteomics Reveals the Dynamics of Proteasome Complexes Composition and Stoichiometry in a Wide Range of Human Cell Lines. *Journal of Proteome Research* **13**, 3027–3037 (2014).

7. M.-P. Bousquet-Dubouch, *et al.*, Affinity Purification Strategy to Capture Human Endogenous Proteasome Complexes Diversity and to Identify Proteasome-interacting Proteins. *Molecular & Cellular Proteomics* **8**, 1150–1164 (2009).
8. D. Bouyssié, *et al.*, Proline: an efficient and user-friendly software suite for large-scale proteomics. *Bioinformatics* **36**, 3148–3155 (2020).
9. J. Lesne, M.-P. Bousquet, J. Marcoux, M. Locard-Paulet, Top-Down and Intact Protein Mass Spectrometry Data Visualization for Proteoform Analysis Using VisioProt-MS. *Bioinform Biol Insights* **13**, 1177932219868223 (2019).
10. M. Gersch, *et al.*, A Mass Spectrometry Platform for a Streamlined Investigation of Proteasome Integrity, Posttranslational Modifications, and Inhibitor Binding. *Chemistry & Biology* **22**, 404–411 (2015).
11. M. Locard-Paulet, *et al.*, VisioProt-MS: interactive 2D maps from intact protein mass spectrometry. *Bioinformatics* **35**, 679–681 (2019).
12. J. Lesne, *et al.*, Conformational maps of human 20S proteasomes reveal PA28- and immuno-dependent inter-ring crosstalks. *Nature Communications* **11**, 6140 (2020).
13. D. Bouyssié, *et al.*, HDX-Viewer: interactive 3D visualization of hydrogen–deuterium exchange data. *Bioinformatics* **35**, 5331–5333 (2019).
14. T. D. Goddard, *et al.*, UCSF ChimeraX: Meeting modern challenges in visualization and analysis. *Protein Sci.* **27**, 14–25 (2018).
15. W. Humphrey, A. Dalke, K. Schulten, VMD: Visual molecular dynamics. *Journal of Molecular Graphics* **14**, 33–38 (1996).
16. B. Webb, A. Sali, Comparative Protein Structure Modeling Using MODELLER. *Curr Protoc Bioinformatics* **54**, 5.6.1-5.6.37 (2016).
17. D. Van Der Spoel, *et al.*, GROMACS: fast, flexible, and free. *J Comput Chem* **26**, 1701–1718 (2005).
18. J. A. Maier, *et al.*, ff14SB: Improving the Accuracy of Protein Side Chain and Backbone Parameters from ff99SB. *J Chem Theory Comput* **11**, 3696–3713 (2015).
19. W. L. Jorgensen, J. Chandrasekhar, J. D. Madura, R. W. Impey, M. L. Klein, Comparison of simple potential functions for simulating liquid water. *Journal of Chemical Physics* **79**, 926–935 (1983).
20. G. Bussi, D. Donadio, M. Parrinello, Canonical sampling through velocity rescaling. *J Chem Phys* **126**, 014101 (2007).
21. M. Parrinello, A. Rahman, Crystal Structure and Pair Potentials: A Molecular-Dynamics Study. *Phys. Rev. Lett.* **45**, 1196–1199 (1980).
22. T. Darden, D. York, L. Pedersen, Particle mesh Ewald: An N·log(N) method for Ewald sums in large systems. *J. Chem. Phys.* **98**, 10089–10092 (1993).
23. U. Essmann, *et al.*, A smooth particle mesh Ewald method. *J. Chem. Phys.* **103**, 8577–8593 (1995).
24. B. Hess, P-LINCS:  A Parallel Linear Constraint Solver for Molecular Simulation. *J. Chem. Theory Comput.* **4**, 116–122 (2008).
25. W. Baumeister, J. Walz, F. Zühl, E. Seemüller, The proteasome: paradigm of a self-compartmentalizing protease. *Cell* **92**, 367–380 (1998).

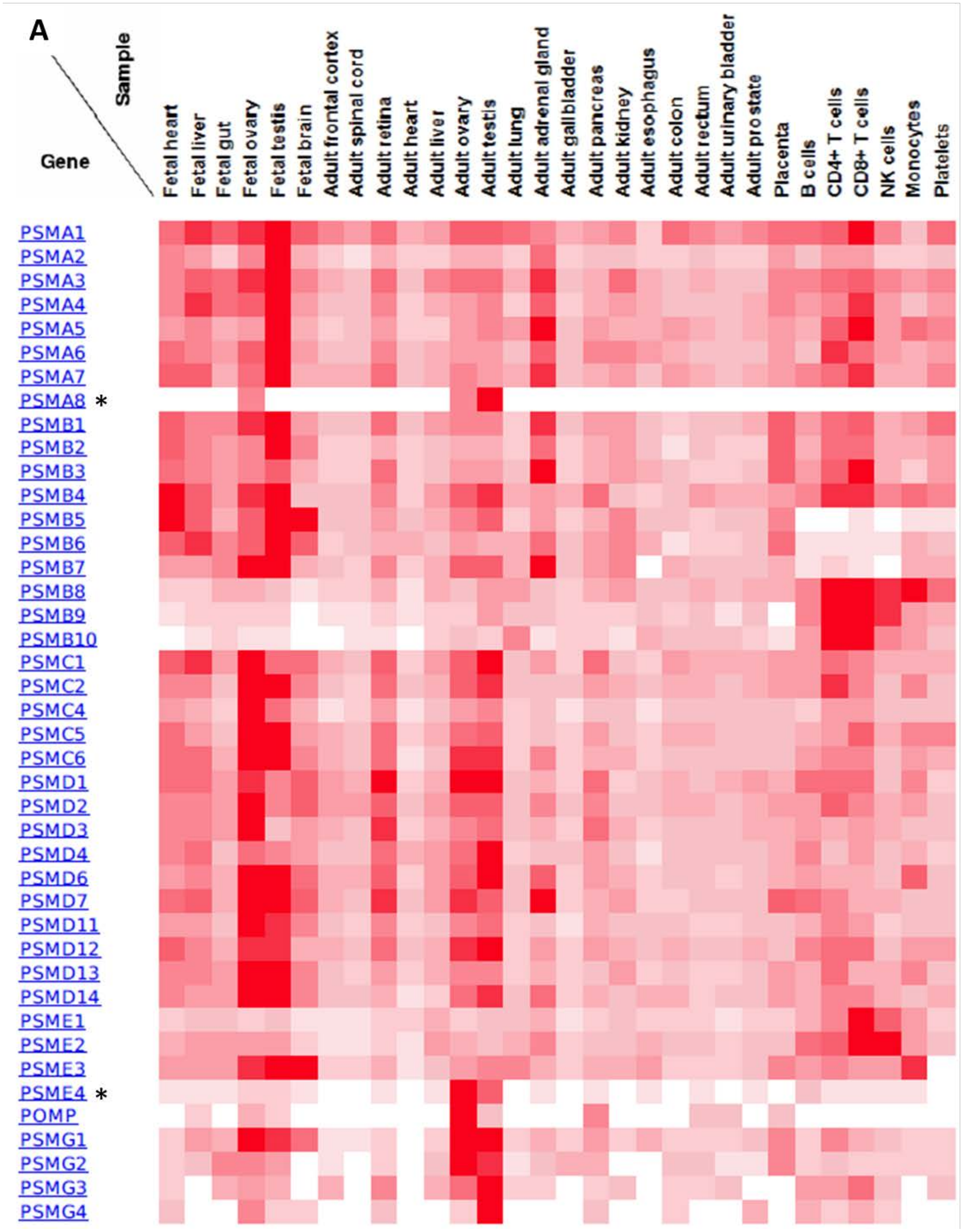

A
Sample
Gene
Fetal heart
Fetal liver
Fetal gut
Fetal ovary
Fetal testis
Fetal brain
Adult frontal cortex
Adult spinal cord
Adult retina
Adult heart
Adult liver
Adult ovary
Adult testis
Adult lung
Adult adrenal gland
Adult gallbladder
Adult pancreas
Adult kidney
Adult esophagus
Adult colon
Adult rectum
Adult urinary bladder
Adult prostate
Placenta
B cells
CD4+ T cells
CD8+ T cells
NK cells
Monocytes
Platelets
PSMA1
PSMA2
PSMA3
PSMA4
PSMA5
PSMA6
PSMA7
PSMA8 *
PSMB1
PSMB2
PSMB3
PSMB4
PSMB5
PSMB6
PSMB7
PSMB8
PSMB9
PSMB10
PSMC1
PSMC2
PSMC4
PSMC5
PSMC6
PSMD1
PSMD2
PSMD3
PSMD4
PSMD6
PSMD7
PSMD11
PSMD12
PSMD13
PSMD14
PSME1
PSME2
PSME3
PSME4 *
POMP
PSMG1
PSMG2
PSMG3
PSMG4

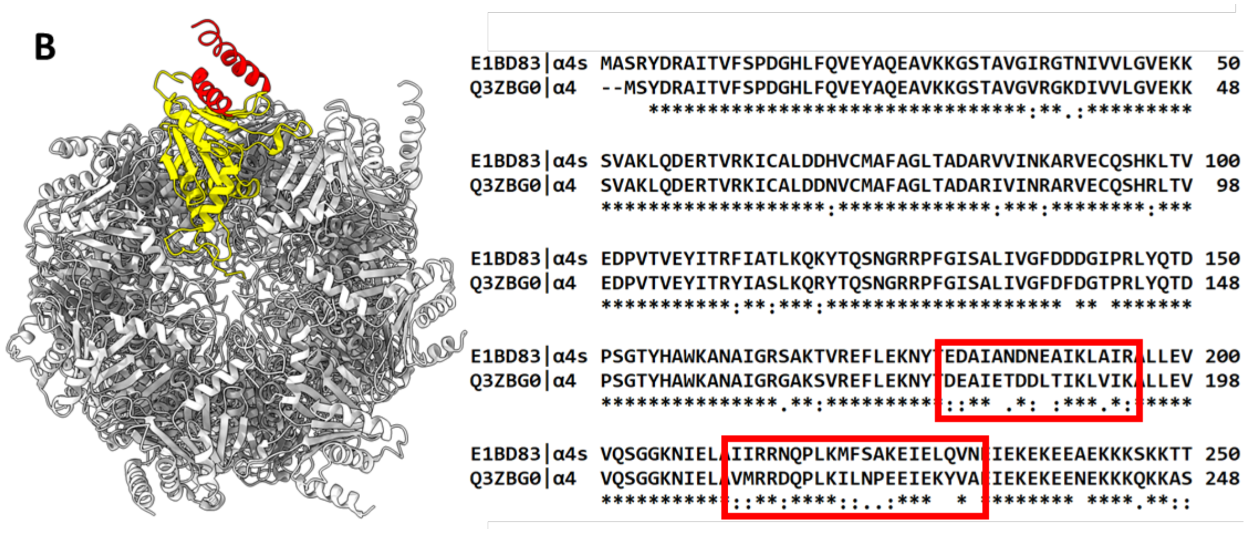


**Figure S1**. **(A)** Expression of all the proteasome subunits and proteasome regulators in different tissues. Of particular interest here are the α4 (PSMA7) subunit and its isoform – α4s (PSMA8) subunit, specific to s20S and proteasome regulators - 19S (PSMC, PSMD), PA28αß (PSME1 & 2), PA28γ (PSME3), and PA200 (PSME4). For the sake of simplicity, gene names are used to represent proteins. Abundances at the protein level are presented across different tissues in a form of a heatmap, generated on Human Proteome Map (Kim *et al.*, 2014). We can see the shared pattern of expression for the PA200 and 19S regulators and the α4s (PSMA8) subunit, suggesting a mutual role. **(B)** Differences between α4 and α4s isoforms. Sequence identity between these two separate isoforms is very high; however, the differences are concentrated in two regions at the C-terminus of only about 30 amino acid residues in total. In α4, this region is completely on the outer surface, exposed to the solvent, and not involved in the α ring interface. Left: α4s (yellow) was modelled on the structure of the c20S. The regions of pronounced differences are marked in red, notably the two outward-facing helices. Right: Sequence alignment of bovine α4 and α4s. The sequences are almost identical, except for the regions marked by the two red squares.

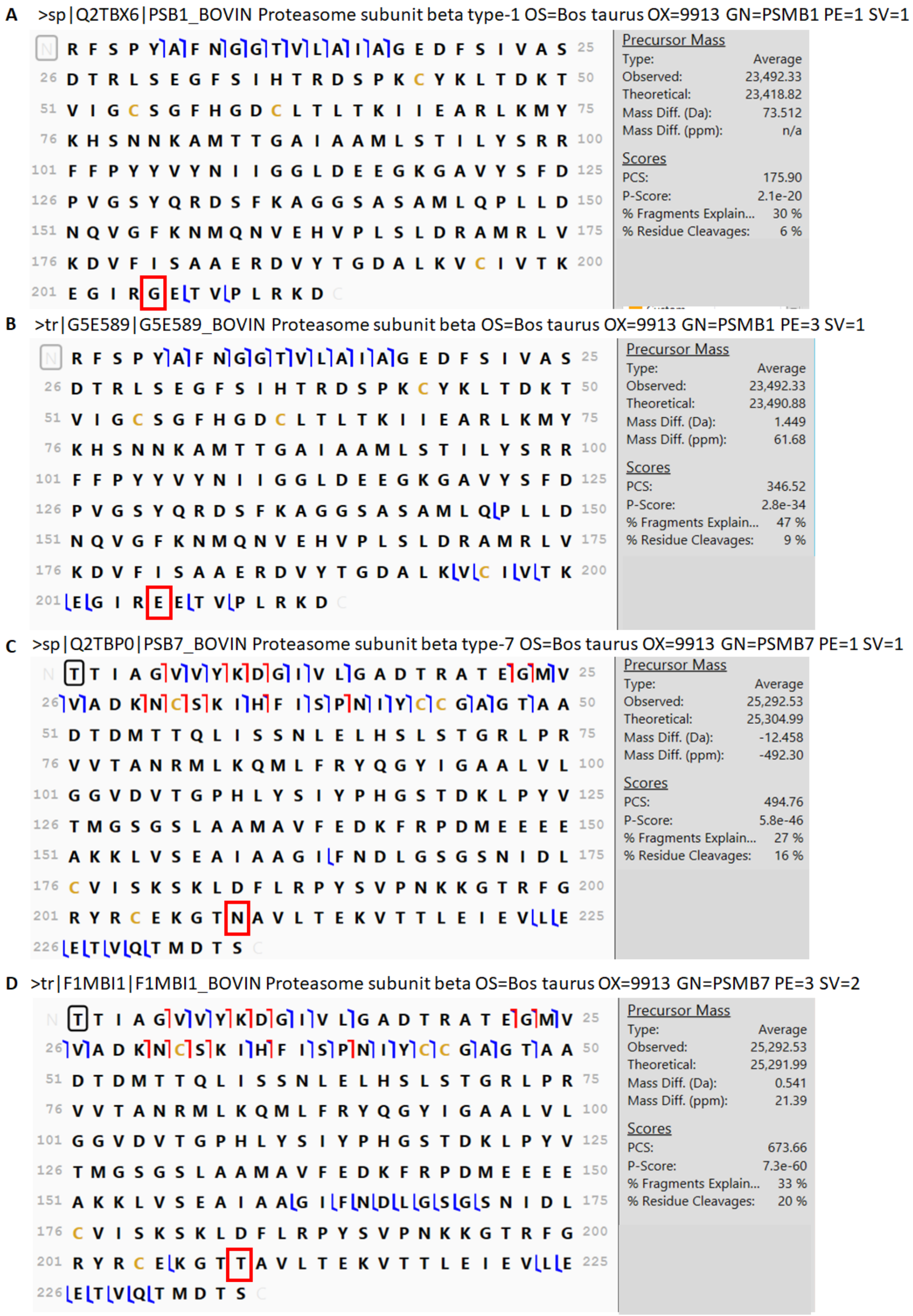


**Figure S2.** The Top-Down analysis of 20S proteasome from bovine testes highlighted errors in the Swiss-Prot entries of **(A)** β6 (Q2TBX6) and **(C)** β2 (Q2TBPO), whereas the TrEMBL counterparts of **(B)** β6 (G5E589: Arg233Glu) and β2 **(D)** F1MBI1 Asn252Thr) provided more fragments and higher identification scores.

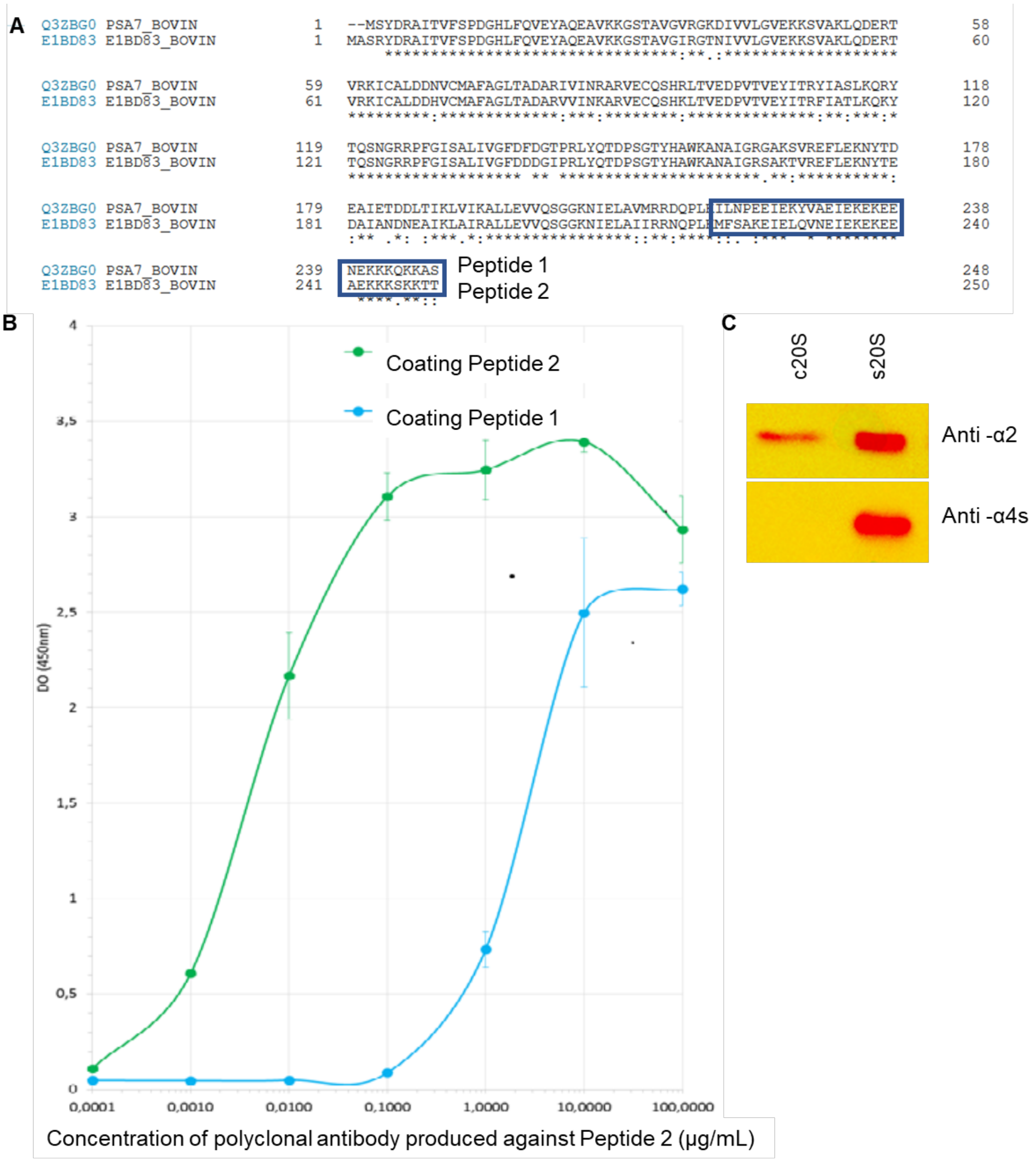


**Figure S3. (A)** Sequence alignment of bovine PSMA7 and PSMA8 proteins showing the C-terminal sequence chosen for the immunization strategy (Peptide 1 : ILNPEEIEKYVAEIEKEKEENEKKKQKKAS and Peptide 2 : MFSAKEIELQVNEIEKEKEEAEKKKSKKTT) **(B)** ELISA assay (plates were coated with peptide 1 (corresponding to the 219-248 C-term part of the α4 protein) or peptide 2 (corresponding to the 221-250 C-term part of the α4s protein) and the different concentration of polyclonal antibody produced against peptide 2 were used for the readout. This resulted in EC50 of 0.005 µg/mL and 2.12 µg/mL for coating peptides 2 and 1, respectively. **(C)** Western-blot showing the specificity of the antibody produced against bovine α4s. c20S proteasome were purified from bovine muscle tissue and s20S (mixed with c20S) was purified from bovine testes. The anti-α2 antibody was obtained from the MCP21 hybridoma.

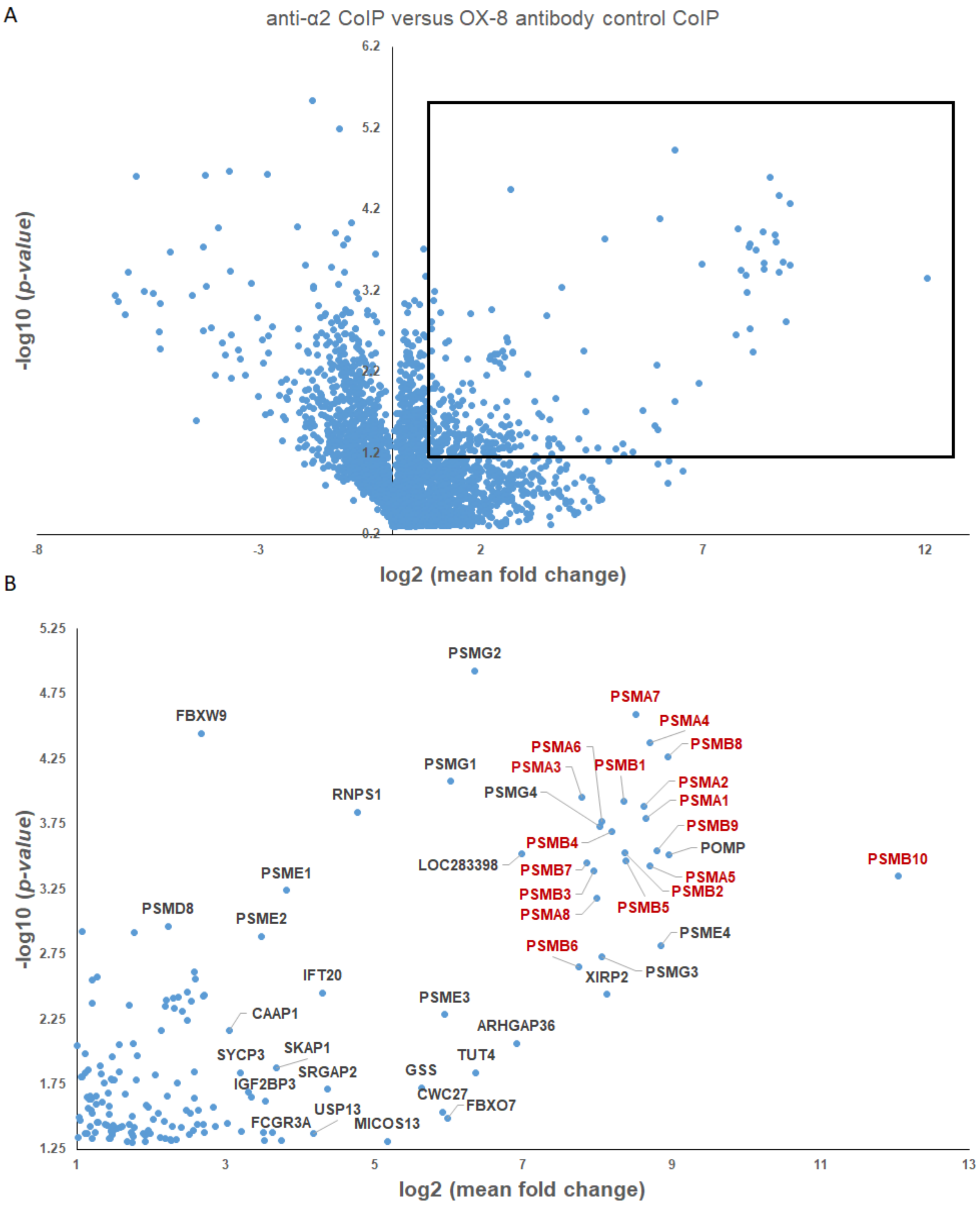


**Figure S4.** Anti-α2 immunopurification **(A)** Volcano plot volcano presenting the statistical significance distribution against the log2-transformed fold change between anti-α2 IP and unrelated control OX8 IP. **(B)** Close-up view of proteins that were significantly enriched (FC>2, *p-value* <0.05) in anti-α2 vs. OX8 IP. For the sake of clarity, gene names are used to represent the proteins.

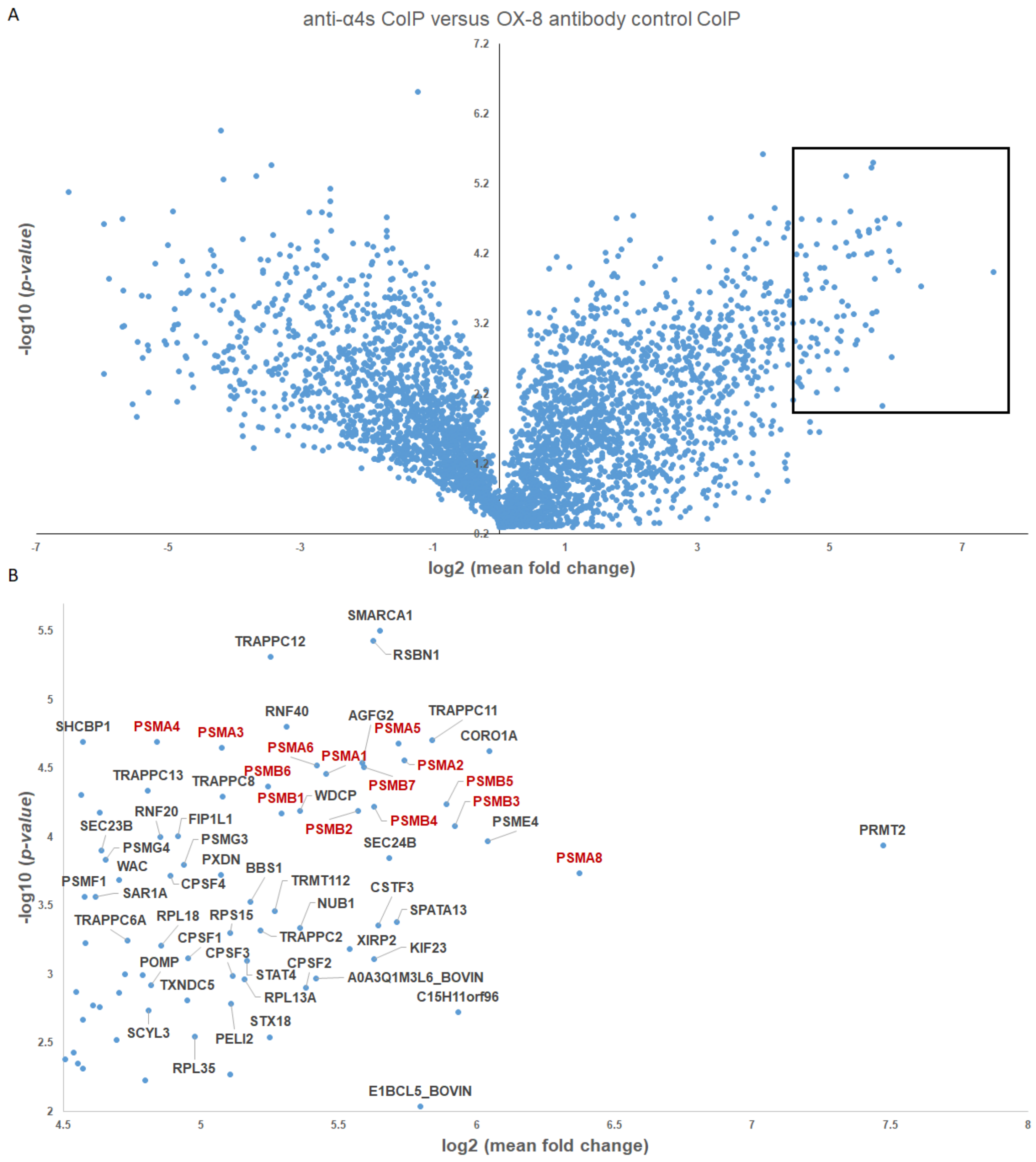


**Figure S5.** Anti-α4s immunopurification **(A)** Volcano plot volcano presenting the statistical significance distribution against the log2-transformed fold change between anti-α4s IP and unrelated control OX8 IP. **(B)** Close-up view of the most significantly enriched proteins in anti-α4s *vs.* OX8 control IP. For the sake of clarity, gene names are used to represent the proteins.

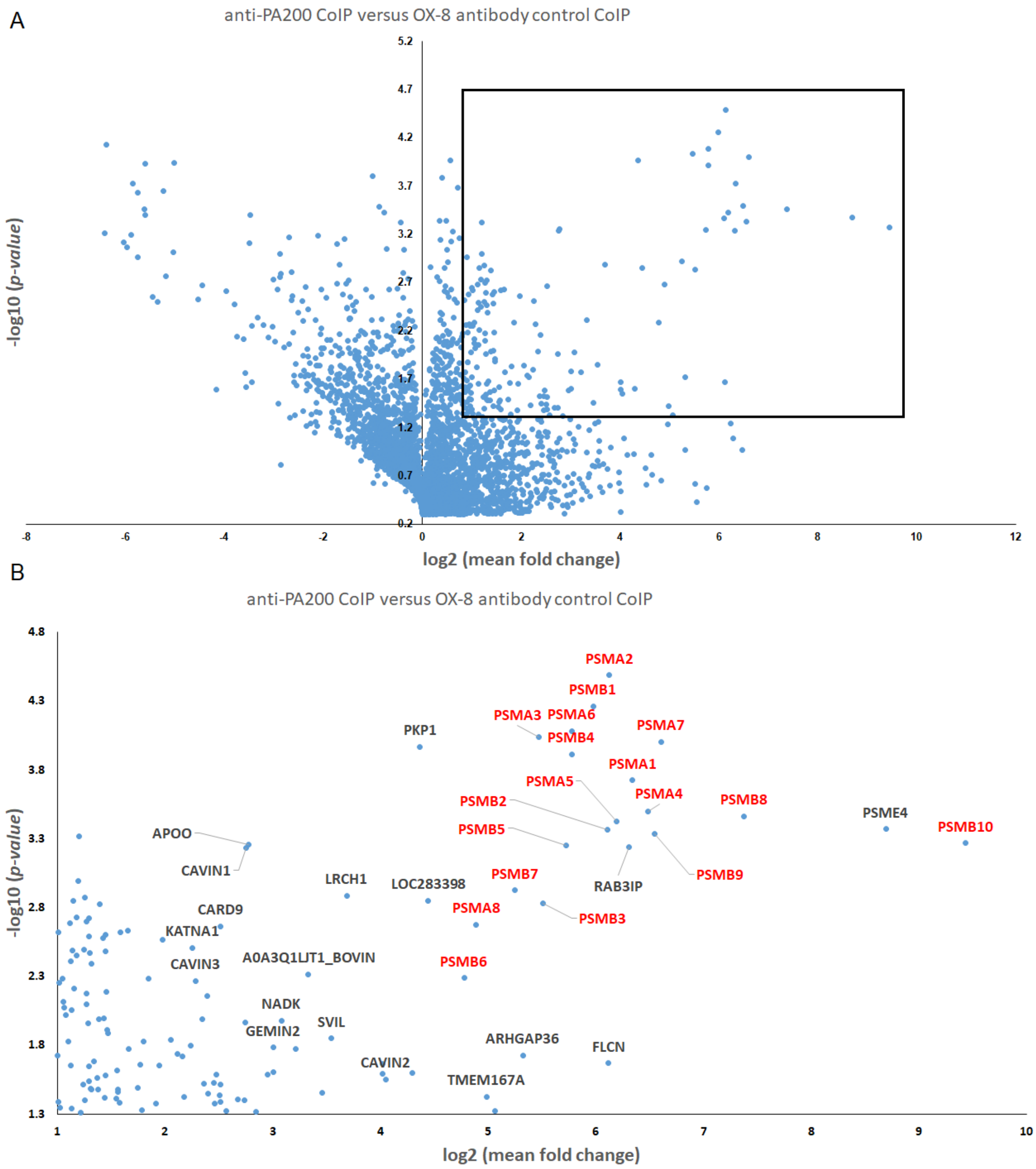


**Figure S6.** Anti-PA200 immunopurification Volcano plot volcano presenting the statistical significance distribution against the log2-transformed fold change between anti-PA200 IP and unrelated control OX8 IP. **(B)** Close-up view of proteins that were significantly enriched (FC>2, *p-value* <0.05) in anti-PA200 vs. OX8 IP. For the sake of clarity, gene names are used to represent the proteins.

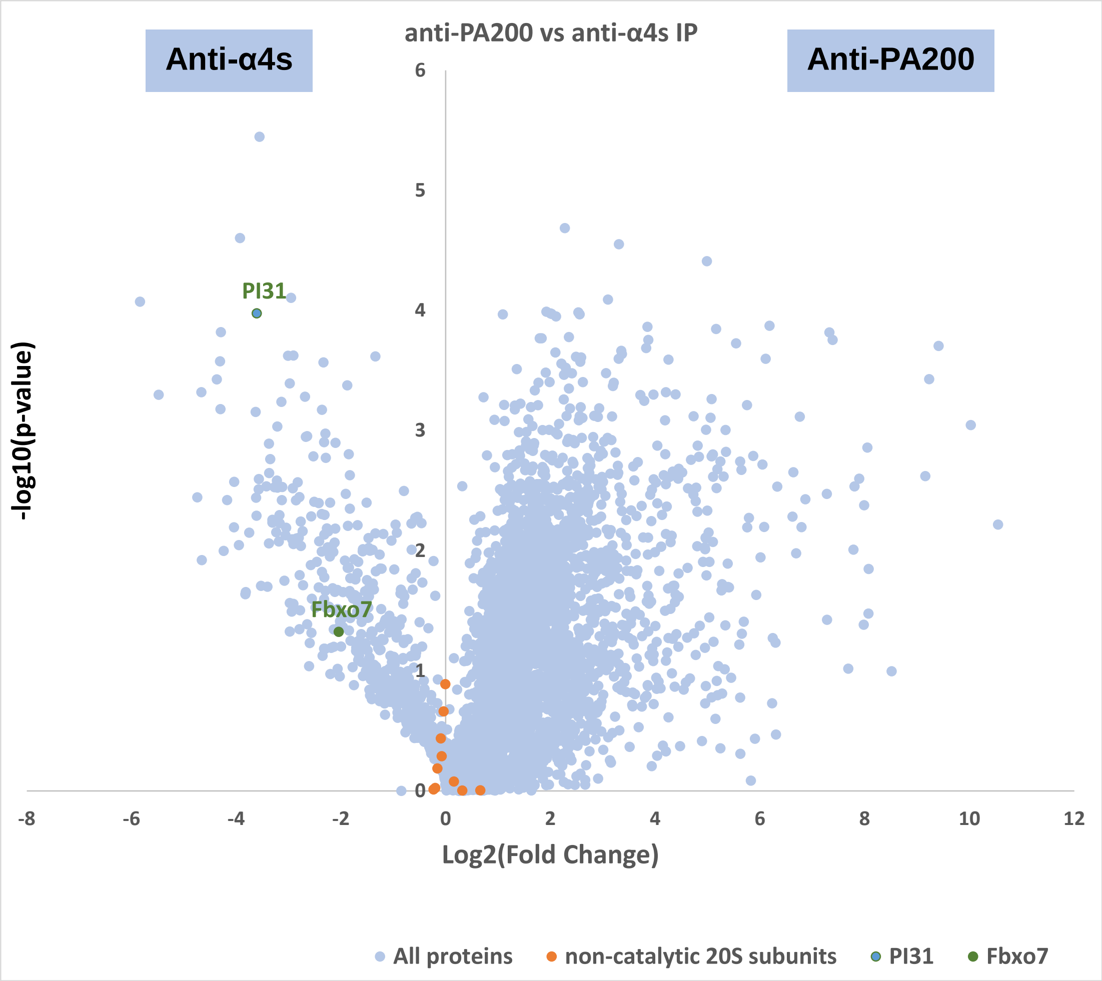


**Figure S7.** Volcano plot volcano presenting the statistical significance distribution against the log2-transformed fold change between anti-α4s (left hand side) and anti-PA200 (right hand side).

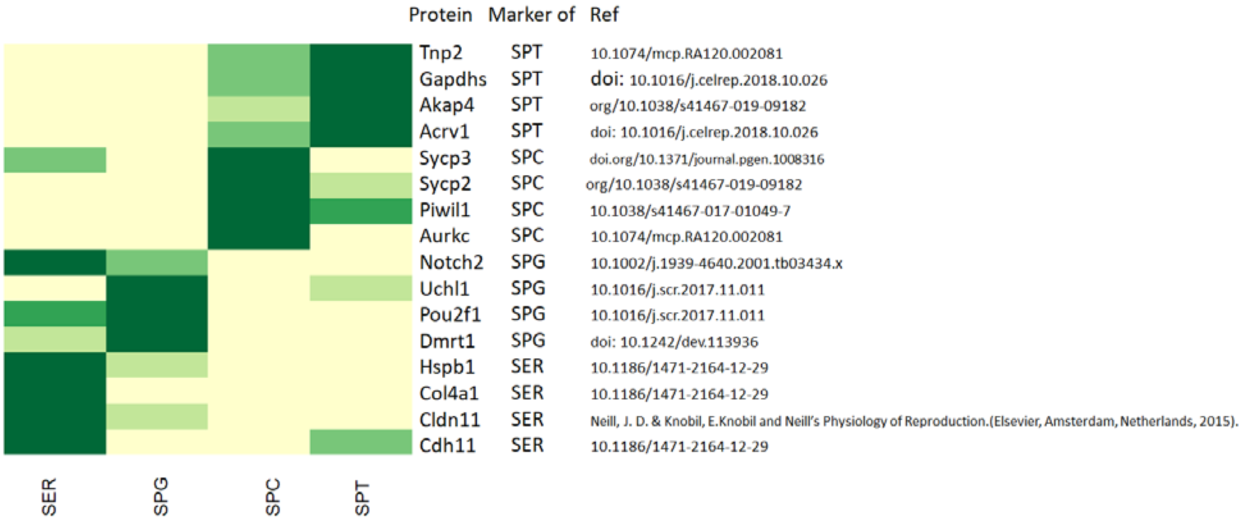


**Figure S8.** Label-free MS quantification of protein markers of sertoli cells (SER), spermatogonia (SPG), spermatocytes (SPC), and spermatids (SPT). Among 5750 proteins validated and quantified in the lysates dataset obtained by LC-MS/MS, a pool of 12 protein markers of each separated cell type were used to validate the cell purification protocol.

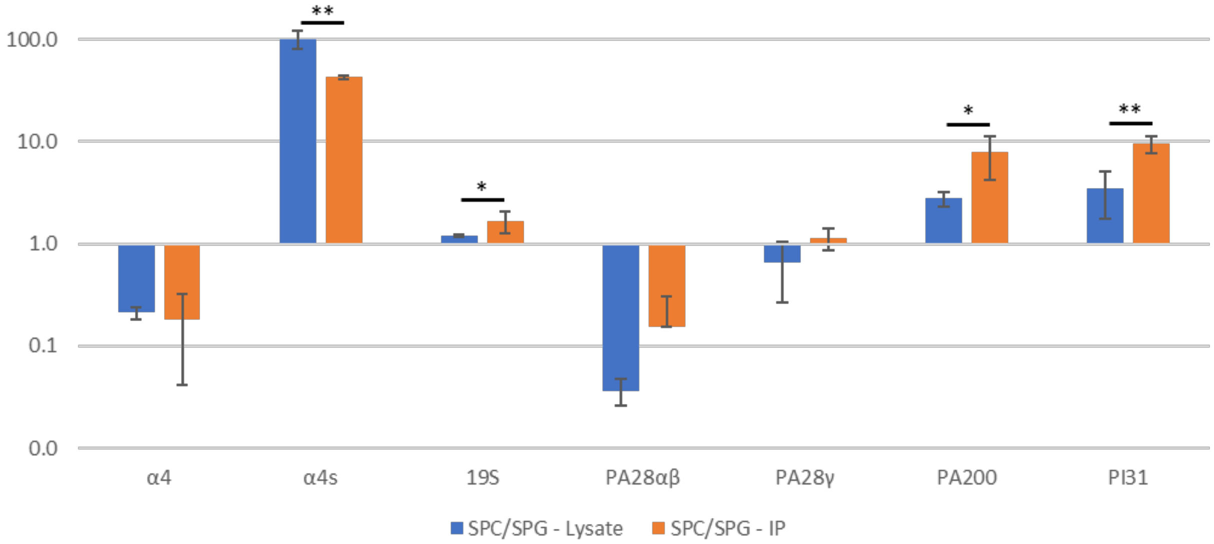


**Figure S9.** Comparison of protein abundances changes from SPG to SPC stages in the lysate and in the immunopurified proteasome. For each protein or protein complex, the change in quantity from SPG to SPC (= ratio SPC/SPG) was calculated both in the lysate and in the immunopurified proteasome sample. Then, these ratios were normalized with the corresponding ratios in the lysates. Stars indicate significance between the lysate and its corresponding immunopurified proteasome sample (IP).

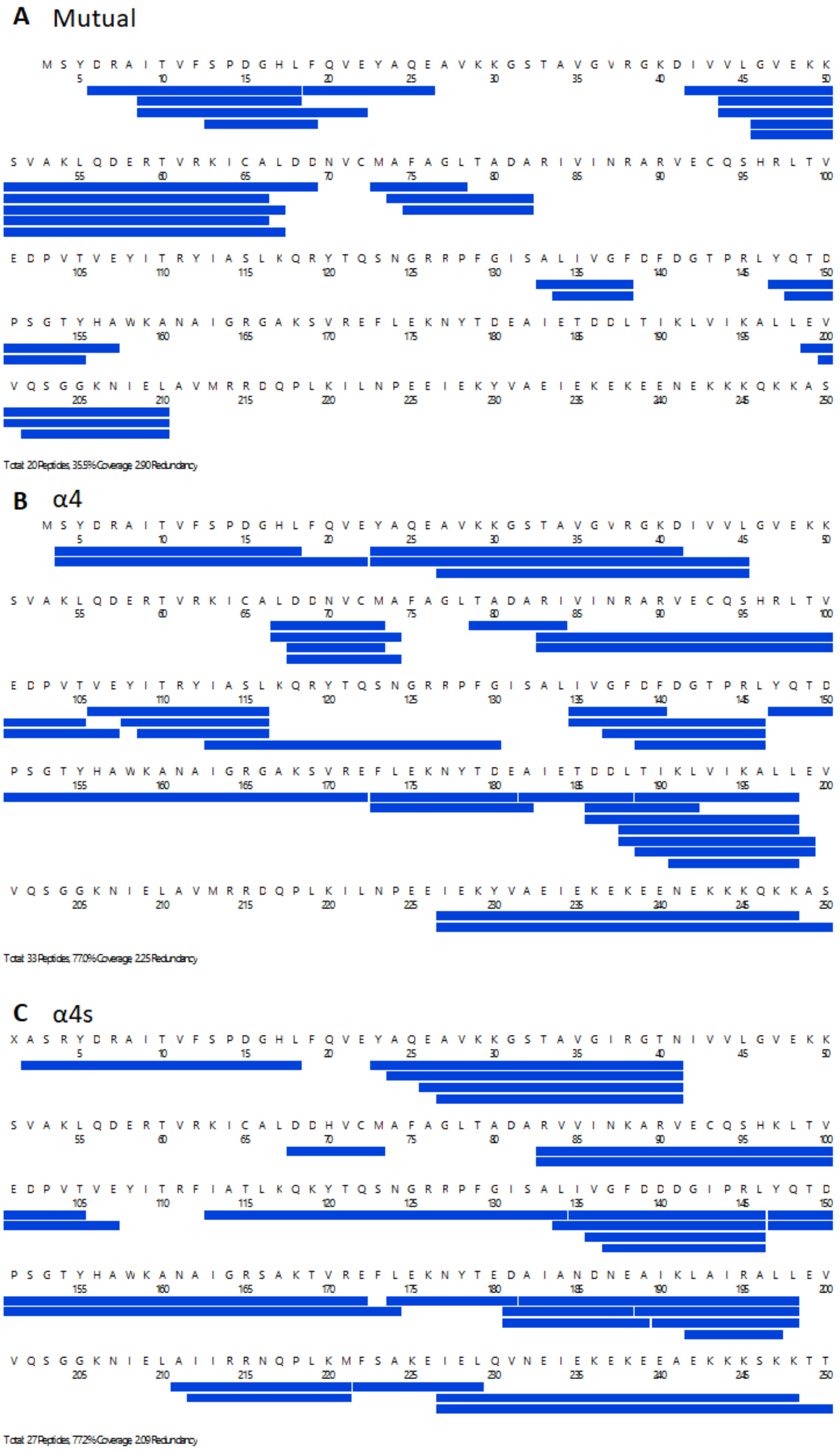


**Figure S10.** Sequence coverage obtained upon pepsin digestion. **(A)** 20 peptides are found in both α4 and α4s. Proteospecific peptides obtained for **(B)** α4 and **(C)** α4s.

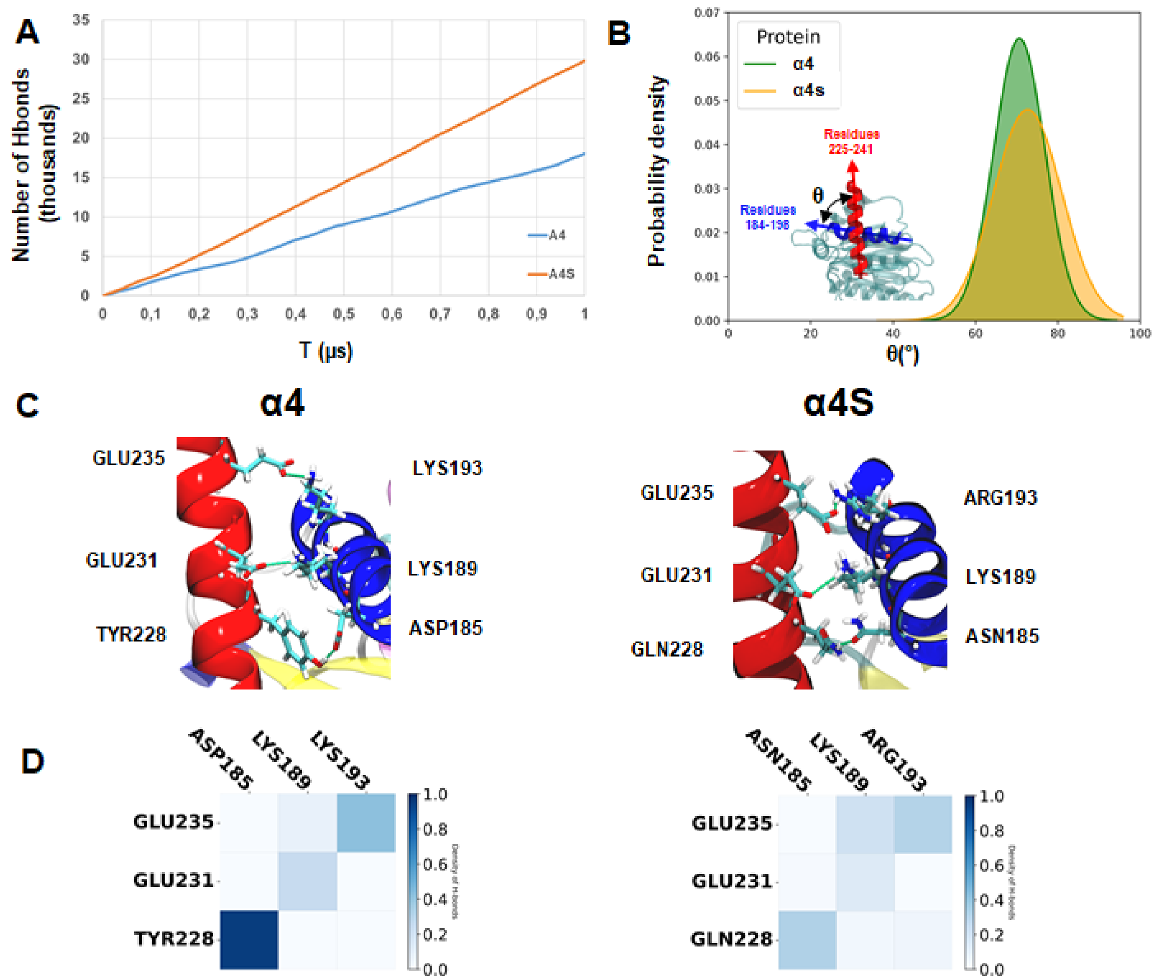


**Figure S11.** Molecular dynamics simulations of α4 and α4s. **(A)** Number of H-bonds observed for the amide protons of residues 184-198 and 225-241 of α4s (orange line) and α4 (blue line) during the 1 μs simulation. **(B)** Distribution of crossing angles between the two last α-helices of α4s (yellow) and α4 (green). **(C)** Close-up view of the two last α-helices of α4 (left) and α4s (right). **(D)** Normalized frequency of H-bonds observed between the side-chain residues of the two last α-helices of α4 (left) and α4s (right).

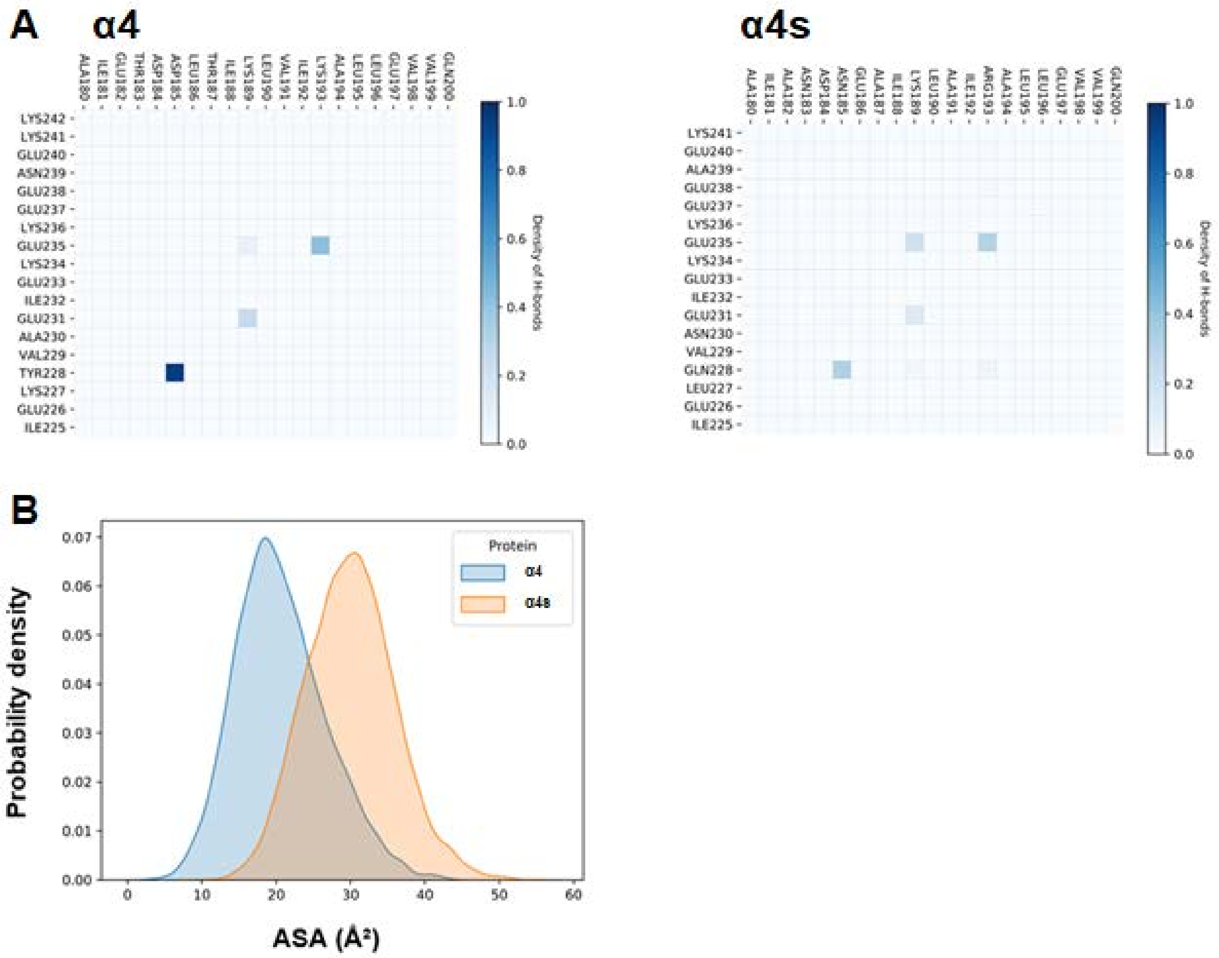


**Figure S12. (A)** Normalized frequency of H-bonds observed between the side-chain residues of the two last α-helices of α4 (left) and α4s (right). **(B)** Distributions of Accessible Surface Areas (Å²) observed for the residues 184-198 and 225-241 α4 (blue) and α4s (red).

| | Run 1 | Run 2 | Run 3 | Average | Standard deviation |
|---|---|---|---|---|---|
| β5i (%) | 35.0 | 36.3 | 36.8 | 36.0 | 0.9 |
| β1i (%) | 32.0 | 29.6 | 30.5 | 30.7 | 1.2 |
| β2i (%) | 17.5 | 7.5 | 9.9 | 11.7 | 5.2 |

**Table S1.** Estimation of the relative abundances of the immuno-catalytic subunits in whole bovine testes (See Top-Down MS experimental part for detailed calculations).

| Name* | Theoretical MW (mature isoform) | Observed mass | Modification | Δm |
|---|---|---|---|---|
| α1 | 27399.5 | 27308.7 | -Met+Ac | -1.8 |
| α2 | 25767.4 | 25807.8 | +Ac | -1.6 |
| α3 | 29483.8 | 29393.2 | -Met+Ac | -1.6 |
| α4 | 27868.9 | 27777.5<br>27620.7 | -Met+Ac<br>-Met+Ac-AS | -2.4<br>-1.0 |
| α4s | 27915.0 | 27825.1<br>27622.5 | -Met+Ac<br>-Met+Ac-TT | -0.9<br>-1.1 |
| α5 | 26411.0 | 26451.9 | +Ac | -1.1 |
| α6 | 29585.6 | 29626.0 | +Ac | -1.6 |
| α7 | 28274.0 | 28394.2<br>28314.9 | +Phos+Ac<br>+Ac | -1.8<br>-1.1 |
| β1 | 22010.1 | 22009.0 | - | -1.1 |
| β1i | 21373.2 | 21371.7 | - | -1.5 |
| β2 | 25292.0 | 25291.3 | - | -0.7 |
| β2i | 24709.3 | 24707.9 | - | -1.4 |
| β3 | 22861.7 | 22902.5 | +Ac | -1.2 |
| β4 | 22896.3 | 22937.1 | +Ac | -1.2 |
| β5 | 22474.3 | 22472.6 | - | -1.7 |
| β5i | 22632.7 | 22649.0 | +Ox | +0.3 |
| β6 | 23490.9 | 23489.6 | - | -1.3 |
| β7 | 24361.8 | 24360.4 | - | -1.4 |

**Table S2.** Theoretical and observed MWs of the bovine proteasome subunits, along with their corresponding PTMs *nomenclature as in (25).

| | α4s/α4 | p-value (T-test) |
|---|---|---|
| non-catalytic 20S | 1.030 | 0.234 |
| catalytic c20S | 3.133 | 0.002 |
| catalytic i20S | 0.006 | 6.8E-05 |
| 19S | 14.555 | 0.025 |
| PA28αβ | 0.164 | 0.012 |
| PA28γ | 0.831 | 0.363 |
| PA200 | 1.969 | 0.0270 |
| PI31 | 1451.3 | 0.002 |
| Fbxo7 | 71.58 | 0.007 |
| PAC1-4 | 0.887 | 0.136 |
| POMP | 0.187 | 0.020 |
| SYCE1 | 294.31 | 0.009 |
| SYCP3 (rep1 excluded) | 125.65 | 0.013 |

**Table S3.** Estimation of the abundance ratio of different proteins between α4s and α4 containing proteasomes in whole bovine testes (See experimental part for detailed calculations).

| Gene name | averaged normalized iBAQ_IP anti-α4s | averaged normalized iBAQ_IP anti-α2 | averaged FC (anti-α4s *vs.* anti-α2) | t test (anti-α4s *vs.* anti-α2) |
|---|---:|---:|---:|---:|
| USP10 | 5.86E-03 | 4.07E-05 | 148.49 | 4.29E-05 |
| RNF40 | 3.15E-02 | 4.51E-04 | 103.05 | 8.83E-04 |
| RNF20 | 3.03E-02 | 4.40E-04 | 84.31 | 4.90E-04 |
| UBE2A | 1.41E-02 | 2.74E-03 | 35.36 | 4.59E-02 |
| DTX3L | 1.80E-03 | 7.59E-05 | 33.26 | 9.08E-03 |
| HUWE1 | 1.94E-03 | 1.10E-04 | 26.55 | 5.39E-03 |
| HERC4 | 7.85E-04 | 7.88E-05 | 19.83 | 1.77E-02 |
| USP16 | 6.28E-04 | 6.48E-05 | 18.89 | 7.17E-03 |
| UFM1 | 1.04E-02 | 5.81E-04 | 18.88 | 4.96E-03 |
| HERC5 | 1.22E-03 | 2.86E-04 | 17.12 | 4.78E-02 |
| USP40 | 2.53E-03 | 3.42E-04 | 16.04 | 1.44E-02 |
| USP15 | 4.96E-04 | 1.10E-04 | 11.45 | 4.71E-02 |
| OTUD4 | 2.04E-04 | 2.52E-05 | 11.45 | 8.78E-03 |
| UFC1 | 1.05E-03 | 9.83E-05 | 10.71 | 5.12E-03 |
| UBR5 | 3.99E-03 | 6.87E-04 | 9.60 | 2.09E-02 |
| OTUB1 | 1.13E-02 | 2.15E-03 | 9.05 | 4.01E-02 |
| UBE3A | 3.40E-03 | 6.20E-04 | 7.98 | 1.11E-02 |
| NPLOC4 | 8.87E-03 | 1.79E-03 | 7.68 | 2.27E-02 |
| USP26 | 2.46E-04 | 3.40E-05 | 7.61 | 1.83E-03 |
| USP34 | 1.81E-04 | 3.68E-05 | 7.57 | 1.92E-02 |
| PELI2 | 8.44E-02 | 1.20E-02 | 7.52 | 3.60E-04 |
| UBR1 | 9.39E-04 | 1.26E-04 | 7.12 | 2.54E-02 |
| USP11 | 1.51E-04 | 2.38E-05 | 7.00 | 3.17E-02 |
| ARIH2 | 2.18E-04 | 4.67E-05 | 6.85 | 4.59E-02 |
| USP5 | 1.58E-02 | 2.30E-03 | 6.66 | 4.05E-04 |
| USP47 | 3.92E-04 | 1.02E-04 | 6.47 | 3.59E-02 |
| USP12 | 5.17E-04 | 9.10E-05 | 6.42 | 2.36E-03 |
| UFD1 | 1.46E-02 | 3.58E-03 | 5.27 | 1.28E-02 |
| RNF114 | 3.42E-04 | 7.91E-05 | 4.56 | 2.37E-03 |

| | | | | |
|---|---|---|---|---|
| UBE4B | 6.28E-05 | 1.61E-05 | 4.51 | 1.55E-02 |
| UBAP2L | 3.25E-03 | 1.33E-03 | 4.49 | 1.73E-02 |
| KCMF1 | 2.61E-04 | 7.17E-05 | 4.45 | 2.17E-02 |
| UBE2N | 7.15E-04 | 2.67E-04 | 4.27 | 4.67E-02 |
| UCHL5 | 9.81E-03 | 3.96E-03 | 3.87 | 4.65E-02 |
| ITCH | 1.00E-04 | 2.66E-05 | 3.48 | 4.48E-03 |
| USP14 | 5.68E-03 | 2.19E-03 | 3.24 | 1.48E-02 |
| BRCC3 | 1.82E-04 | 7.70E-05 | 2.54 | 4.33E-02 |
| RPS27A | 4.98E-02 | 2.88E-02 | 1.89 | 3.09E-02 |
| ADRM1 | 2.19E-02 | 1.22E-02 | 1.83 | 1.04E-02 |
| HERC2 | 2.55E-06 | 5.69E-06 | 0.49 | 2.82E-02 |
| TRIM21 | 2.68E-03 | 6.71E-02 | 0.04 | 3.58E-05 |

**Table S4.** Averaged normalized IBAQ values and FC enrichment in anti-α4s *vs.* anti α2 IP for selected ubiquitin-related proteins.
